# Machine learning-based structural classification of lytic polysaccharide monooxygenases

**DOI:** 10.1101/2025.07.02.662747

**Authors:** Aathi Manikandan, Ragothaman M. Yennamalli

## Abstract

Lytic polysaccharide monooxygenase (LPMO) is a copper-dependent redox enzyme and according to CAZy is classified either as cellulolytic or chitinolytic. According to CAZy, there are eight families of LPMO namely AA9, AA10, AA11, AA13, AA14, AA15, AA16, and AA17, where AA stands for Auxiliary Activity. Previously, using the sequence information machine learning-based functional annotation was successfully completed using neural network and LSTM models. This was done for AA9 and AA10 as the number of sequences was large enough to train a model with high-performance indices. Here, the goal is to use existing 3D structures and AI-based models of the remaining LPMO sequences and train a machine learning model to identify and classify LPMO with the help of structural features by performing either a neural network algorithm or other suitable methods. Using the features of LPMO such as surface depth, accessible area, electrostatic charge distribution, and geometric features (independent features that define the shape and are not based on enzyme reaction mechanism) we will identify the features with high signal-to-noise ratio/significance using ensemble feature selection method. The features were extracted using Pymol, MdTraj, Biopython, Open3d, and Bio3D tools from OBJ and STL format. The model thus trained using structural features will enable identifying and annotating newer LPMO sequences belonging to one of the eight AA families.

## Introduction

Glycoside hydrolases (GHs) are enzymes that hydrolyze glycosidic bonds in carbohydrates, including polysaccharides, oligosaccharides, and disaccharides, playing key roles in digestion, cellular signaling, and plant biomass degradation (Wang et al., 2021). They use water to cleave these bonds, releasing monosaccharides or smaller oligosaccharides, though the process can be complex and slow (Vaaje-Kolstad et al., 2005). The identification of Lytic polysaccharide monooxygenases (LPMOs) in 2010 transformed GH efficiency. LPMOs employ oxygen and a copper cofactor to oxidize polysaccharides, weakening glycosidic bonds and enhancing cleavage by GHs. This innovation is crucial in industries like biofuel production and waste recycling (Arora et al., 2019).

Lytic polysaccharide monooxygenases (LPMOs), classified as auxiliary activity (AA) enzymes, are efficient catalysts for degrading marine and terrestrial biomasses (Eibinger et al., 2014; Harris et al., 2014). Polysaccharides like hemicellulose, cellulose, chitin, and starch are common components of these biomasses (Wang et al., 2021). Biomass, unlike other renewable resources, can be rapidly converted into liquid biofuels such as ethanol, biodiesel, and green diesel, offering sustainable alternatives to non-renewable fuels. Industrial biotechnology plays a crucial role in transitioning from polluting fossil fuels to carbon-neutral energy sources. However, breaking down recalcitrant polysaccharides like crystalline cellulose remains a significant challenge. Cleaving the glycosidic bond between monomers a critical step in biofuel preparation is facilitated by LPMOs and glycoside hydrolases.

LPMOs were first identified in 1992, but their distinct functionality was revealed only in 2010 (Vaaje-Kolstad et al., 2010). Initially classified as CBM33 in bacteria and GH61 in fungi, LPMOs are now categorized under AA families in the Carbohydrate Active Enzymes (CAZy) database (Aachmann et al., 2015; Levasseur et al., 2013). Eight AA families (AA9-AA17) are recognized, with bacterial and viral enzymes predominantly classified as AA10, and fungal enzymes as AA9, AA11, and AA13. LPMOs are further categorized by substrate specificity: C1-binding (LPMO1), C4-binding (LPMO2), and dual C1/C4-binding (LPMO3) types (Arora et al., 2019).

LPMOs have been identified across bacteria, fungi, viruses, and eukaryotes. Research continues to uncover new sequences, supported by semi-automated and manually curated updates in the CAZy database (Lombard et al., 2013). Despite these advancements, functional annotation of LPMOs based on structure remains underexplored, and this study aims to address this gap.

AA9 and AA10 are the major LPMO families. AA9 includes cellulose-active enzymes previously classified as GH61, while AA10 comprises CBM33 enzymes, originally considered chitin-specific. Interestingly, despite low sequence similarity (20–30%), structural similarities among LPMOs acting on cellulose and chitin reach 80–90% (Zhou, X et al., 2019). Earlier attempts to annotate LPMOs using long short-term memory (LSTM) neural networks highlighted the limitations of sequence-based classification due to low similarity (Srivastava et al., 2020).

Given these challenges, structural analysis provides a more reliable basis for LPMO classification. Structural features, especially differences in catalytic sites, are critical for distinguishing cellulose- and chitin-specific LPMOs, making structural annotation a promising approach for predicting biological activity.

Inspired by Machine Learning (ML) models for automated analysis, we developed a workflow to train models using structural features. A recent study by Dai et al. (2021) demonstrated an ML approach for optimizing aircraft design by analyzing gaps and flushes between frames using 3D structural data. They extracted seam features from raw 3D geometry, identifying critical distances that improved aerodynamic performance. Feature extraction from 3D models can be approached through content-based methods, 3D scene registration, or object recognition (Yu, F et al., 2010).

Similarly, in biology, the RCSB Protein Data Bank (PDB) provides 3D structural data for proteins, stored in files containing atomic coordinates and other molecular information. These raw PDB files can be converted into 3D models using tools like PyMOL. While the CAZy database primarily hosts sequence data, advances in next-generation sequencing (NGS) have significantly expanded its content. Protein sequences from CAZy can be modeled into 3D structures using tools like AlphaFold2.

This study explores using protein structural data to train ML models capable of annotating newly identified proteins. By converting PDB files into 3D models, we extract structural features to differentiate chitinolytic and cellulolytic LPMOs. Our goal is to develop an optimized model that predicts protein function based on structural features, advancing functional annotation with 3D structural information.

## Results and Discussion

LPMOs are classified into eight Auxiliary Activity (AA) families, encompassing both chitinolytic and cellulolytic enzymes. While structural similarities within families exceed 80%, traditional structural classification methods often lack sensitivity due to low sequence similarity within the same family. This prompted earlier efforts to annotate LPMOs using sequence data, with AA9 and AA10 being the most extensively studied families due to their abundance of protein structures and sequences.

The CAZy database provides LPMO-related data, including family names, organism types, scientific names, and GenBank accession IDs. Among the eight families, 14,099 sequences and 167 experimentally determined structures are available. Using GenBank IDs, we identified 6,507 annotated sequences (cellulolytic or chitinolytic) and 7,592 unannotated sequences. Annotated sequences were curated to isolate LPMO domains, excluding regions with unrelated domains. These curated sequences were modeled using AlphaFold2, a deep- learning tool for predicting protein structures (Jumper et al., 2021; Mirdita et al., 2022).

For model preparation and evaluation, experimentally determined LPMO structures were downloaded from CAZy, cross-verified with UniProt, and cleaned of non-protein entities. Structures capable of depolymerizing both cellulose and chitin were excluded. After preprocessing, 146 structures remained: 100 AA9 cellulolytic LPMOs and 46 AA10 chitinolytic LPMOs. Other LPMO families were excluded due to either dual activity or lack of functional annotations related to cellulose or chitin binding. An additional 22 structures were discarded due to missing annotations, leaving 100 cellulolytic and 46 chitinolytic LPMOs. These remaining structures were further modeled using AlphaFold2, and pLDDT scores (Supplementary Figure 1) were calculated to assess structural quality using in-house Python scripts.

To address potential data skewness in chitinolytic LPMOs, we used the structure 2BEM as a query in DALI. This analysis identified 49 high-similarity structures (Z-scores ranging from 10.5 to 5.1), which were confirmed to be distinct from the 100 cellulolytic structures. This final dataset ensures robust input for downstream analyses and model development.

To develop a binary classification model for distinguishing between chitinolytic and cellulolytic LPMOs, we utilized features generated from structural data. A 60:40 ratio was applied for splitting the dataset into training and test sets. To prevent biases or performance issues associated with imbalanced datasets, the training set maintained a near 1:1 ratio of positive (chitinolytic) to negative (cellulolytic) samples. This balanced approach addresses concerns raised in prior studies (Liu. H et al., 2018; Liu. S et al., 2018) about the negative impact of skewed datasets on machine learning models.

In total, 195 structures were included in the dataset, evenly divided between chitinolytic and cellulolytic LPMOs. This ensured that both classes were equally represented, optimizing the model’s predictive performance.

## Feature generation

For feature extraction of 3D topological properties, protein PDB files were converted into "Standard Triangle Language" (STL) format using an in-house Python script executed in PyMOL (Supplementary Figure 2). The STL files were then processed to extract features. The Molecular Surface Package (MSMS) was used to compute reduced surface properties by converting PDB files into XYZ atom coordinate files. Additional features were extracted using in-house Python scripts, and all features were stored in a comma-separated value (CSV) file for analysis. A total of 34 distinct features were generated from MSMS and in- house code. To identify the most significant features for training the machine learning model, we employed Ensemble Feature Selection (EFS) software (http://efs.heiderlab.de/), which combines multiple feature selection methods to generate normalized ensemble scores (Neumann et al., 2017). Features with ensemble scores above 0.5 were considered significant.

From the analysis (Table 1), we identified Convexity, Analytical Surface Area - Solvent Accessible Surface (ASA_SAS), Analytical Surface Area – Solvent Excluded Surface (ASA_SES), Analytical Surface Area – reent (ASA_reent), and Numerical volumes and area – Solvent Excluded Surface Area (NVA_SES_area) as significant features for training. These features, detailed further in Sanner et al. (1996), will serve as critical inputs for the machine learning model.

**Table 1:** Ensemble scores for the extracted features of the LPMO dataset. We observed significant features of Analytical Surface Area - Solvent Accessible Surface (ASA_SAS), Analytical Surface Area – Solvent Excluded Surface (ASA_SES), Analytical Surface Area -(ASA_reent), and Numerical volumes and area – Solvent Excluded Surface Area (NVA_SES_area) for model preparation with an ensemble score of more than 0.6

| Feature | Ensemble score | Feature | Ensemble score |
| --- | --- | --- | --- |
| Convexity | 0.671576 | ASES_R_h | 0.199464 |
| ASA_SAS | 0.539669 | T_faces | 0.186522 |
| ASA_SES | 0.532397 | T_edges | 0.181884 |
| ASA_reent | 0.523722 | T_vetices | 0.181688 |
| NVA_SES_area | 0.519863 | ASES_genus | 0.176169 |
| Concavity | 0.496177 | T_genus | 0.171852 |
| ASA_toric | 0.470544 | RS_genus | 0.166285 |
| NVA_SES_volume | 0.464061 | RS_faces | 0.159101 |
| Convex_volume | 0.453169 | RS_edges | 0.158257 |
| Mesh_volume | 0.423608 | ASES_fac | 0.147782 |
| ASA_contact | 0.417479 | ASES_edg | 0.141286 |
| Compactness | 0.417458 | RS_vertices | 0.140034 |
| Surface_Area | 0.417277 | ASES_vert | 0.139027 |
| Scale | 0.407607 | RS_free_edges | 0.132444 |
| Sphericity | 0.401792 | ASES_s_vert | 0.057093 |
| Asphericity | 0.295058 | T_density | 0.054503 |
| ASES_C_h | 0.233338 | ASES_s_edg | 0.05261 |

### Model generation and Optimization

We implemented several feature-based machine learning algorithms to distinguish between chitinolytic and cellulolytic LPMOs. These included Logistic Regression with solvers (lbfgs, liblinear, newton-cg, newton-Cholesky, sag, saga), Naive Bayes, Random Forest, Support Vector Machine (SVM) with various kernels (linear, polynomial, RBF, sigmoid), and Multi- Layer Perceptrons (MLP) using solvers (adam, lbfgs, sgd) and activation functions (identity, logistic, relu, tanh). All models were developed using in-house Python scripts. While these methods effectively classify proteins, a limitation arises from potential information loss during feature generation. SVM, in particular, is well-suited for classification tasks and has demonstrated the ability to classify protein structures with up to 90 % structural similarity (Cai et al., 2003). To identify the best-fit model, various algorithms and parameter combinations were tested. For further details on machine learning algorithms, see Pedregosa et al. (2011).

After initial implementation, each ML script was modified to dynamically adjust algorithm parameters through iterative loops, creating multiple models with varying configurations. This approach enabled the evaluation of different parameter sets, leading to models with varied performance.

F1-score was used as the primary criterion for optimization. The F1-scores of all models were stored in matrices using NumPy, a Python module. These matrices were analyzed to identify the parameter values that optimized model performance. This systematic parameter tuning ensured the selection of the most effective model for distinguishing between chitinolytic and cellulolytic LPMOs.

### Logistic Regression

We used the logistic regression algorithm from the scikit-learn Python module. Six different solvers in logistic regression were evaluated to plot F1 score graphs for finding the best-fit model (Supplementary Figure 1). The logistic regression model, after tuning hyperparameters, achieved maximum F1 scores of 0.62, 0.62, 0.62, 0.32, and 0.32 using the *lbfgs*, *newton-cg*, *newton-cholesky*, *sag*, and *saga* solvers, respectively (Supplementary Figure 1). Even though there is a slight difference in the performance of the models on the validation set, the differences are not significant when evaluated on the independent set. The F1 scores were 0.65 for the *lbfgs* and *liblinear* solvers, 0.63 for the *newton-cg* and *newton-cholesky* solvers, and 0.62 for the *sag* and *saga* solvers (Supplementary Table 2). The best model using liblinear had a F1 score, precision and recall of 0.64 with an AUC of 0.66 on the ROC curve (Supplementary Figure 2).

### Naïve Bayes

We applied Gaussian Naïve Bayes for classification, achieving an F1 score, precision, and recall of 0.96, with an AUC of 0.92 on the ROC curve (Supplementary Figure 3). The confusion matrix showed 38 (TP), 2 (FP), 1 (FN), and 37 (TN) as shown in Table 2. However, for the independent dataset Gaussian Naïve Bayes performed very poorly with zero TP, 1472 FP, 0 FN, and 517 TN. Also, it had a recall of 0.26, precision of 0.07, and an F1 score value of 0.11 (Table 2). While the validation set gave better performance metrics, the trained model using Gaussian Naïve Bayes gave significantly lower performance metrics.

**Table 2:**
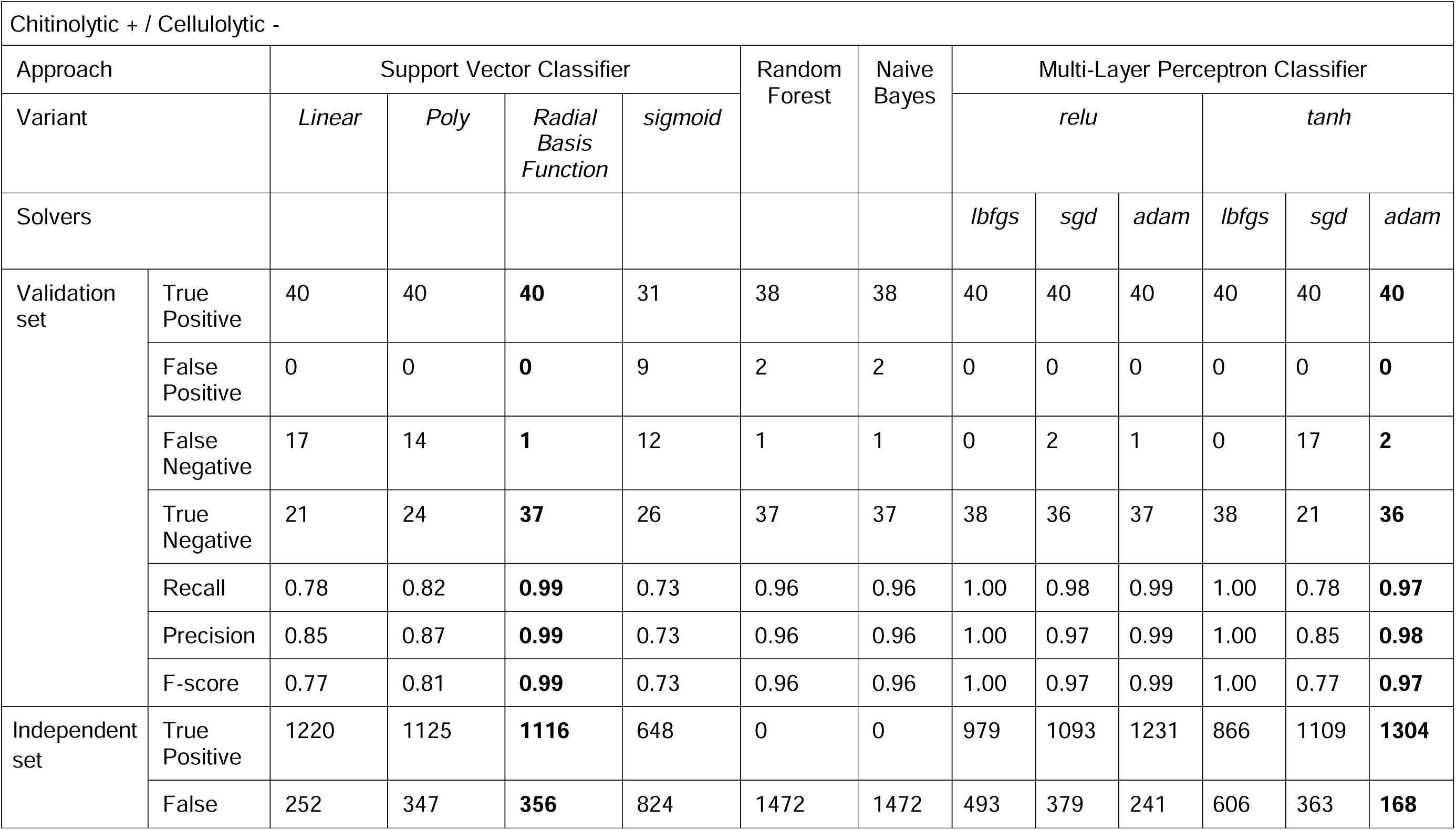
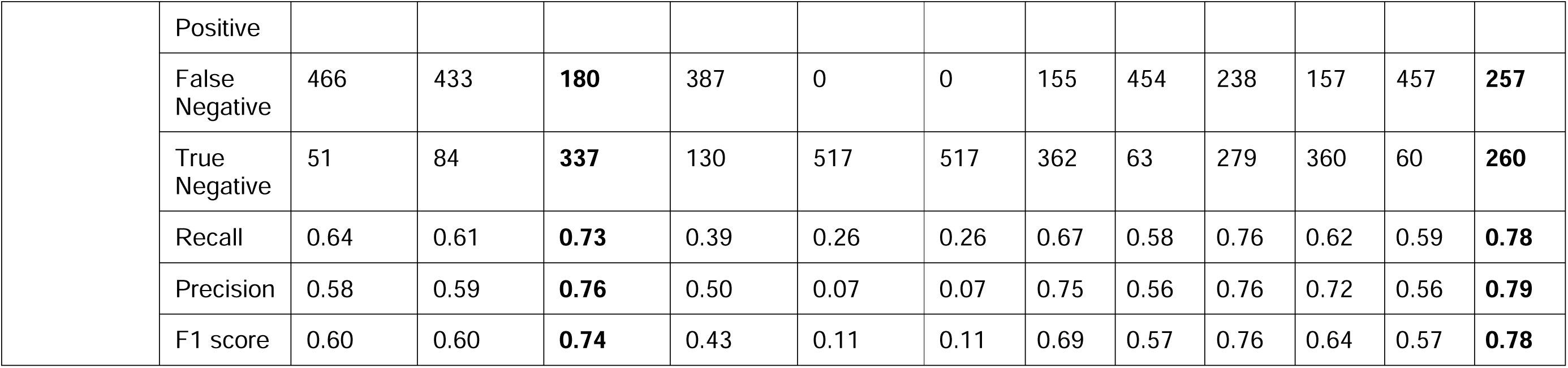
Performance Comparison of Machine Learning Classifiers for Predicting Chitinolytic-Positive / Cellulolytic-Negative Enzymes on Validation and Independent Datasets

### Random Forest

A model was developed using the Random Forest algorithm. The same modifications were applied to the script to generate an F1 score plot for identifying the optimal parameter. As shown in Supplementary Figure 4, the Random Forest model achieved the highest F1 score of 0.96 when the n_estimators parameter reached a value of 15 or higher. The confusion matrix for the validation set consisted of 38 true positives (TP), 2 false positives (FP), 1 false negative (FN), and 37 true negatives (TN), as presented in Table 2. The model achieved a precision of 0.96, a recall of 0.96, and an AUC of 1.00. However, the model did not generalize well to the independent dataset. The corresponding confusion matrix showed 0 TP, 1472 FP, 0 FN, and 517 TN, as shown in Table 2. This indicates that the model overfitted to the validation data and failed to replicate its performance on unseen data, resulting in a significantly lower F1 score of 0.11 (Supplementary Figure 5).

### Support Vector Machine

After evaluating the SVM model with Linear, Polynomial, Radial Basis Function (RBF), and Sigmoid kernels, the results were compared. The model achieved its highest F1 score of 0.99 using the Radial Basis Function (RBF) kernel as shown in Figure 1. The SVM model after optimization using linear kernel gave a F1 score of 0.77 (Supplementary Figure 3), using polynomial kernel gave a F1 score of 0.81 (Supplementary Figure 4), using sigmoid kernel gave a F1 score of 0.73 (Supplementary Figure 5). The optimal parameters were found to be C=2, which helped minimize false negatives, and γ\gamma = 4.001, resulting in an accuracy of 98.72%. The confusion matrix for the validation set consisted of 40 true positives (TP), 0 false positives (FP), 1 false negative (FN), and 37 true negatives (TN). The model achieved a precision and recall of 0.99 each, with an AUC score of 1.00 as observed in the ROC curve (Figure 2). Furthermore, the model demonstrated strong performance on the independent dataset, with a confusion matrix of 1096 TP, 376 FP, 170 FN, and 347 TN, as shown in Table 2. The F1 score for the independent set was 0.74, the highest among the various SVM model variants evaluated.

**Figure 1:**
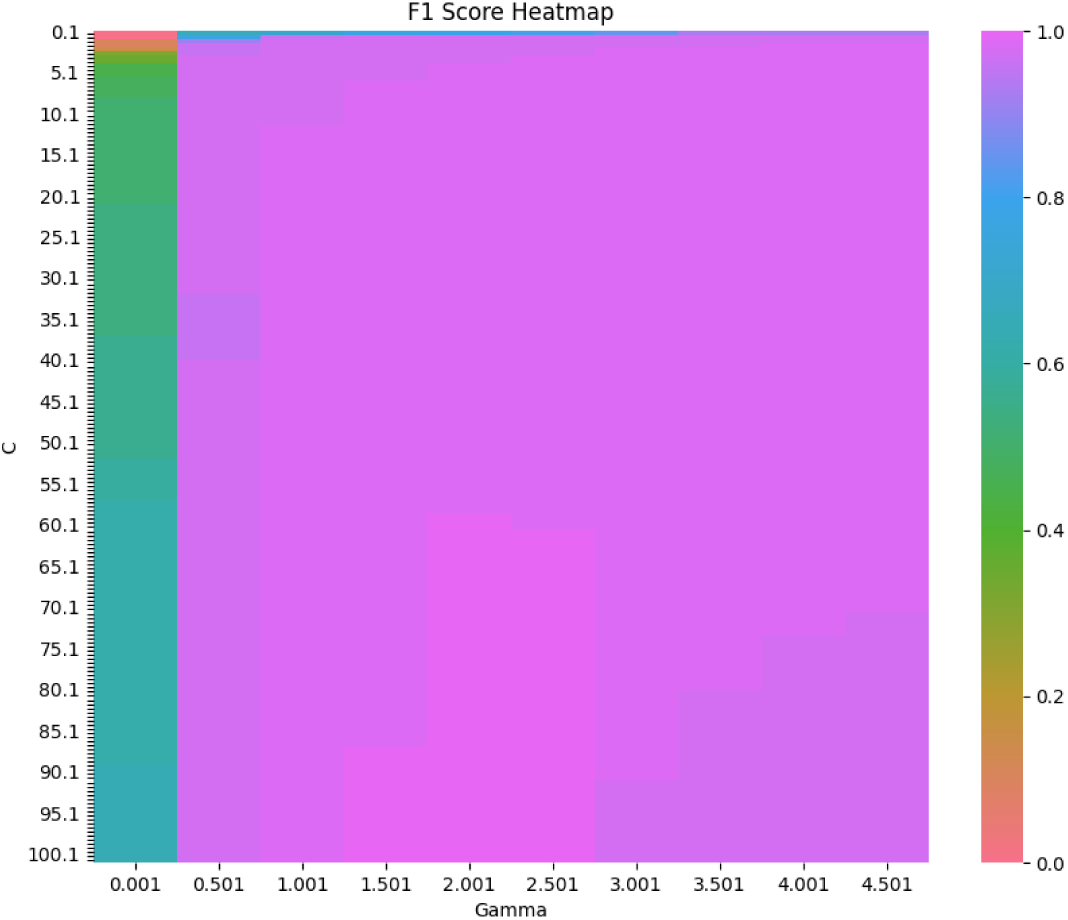
Heatmap plot of F1 score values for Radial Basis Function kernel in SVM with a maximum F1 score of 0.99 is used for the best model. The colors indicate the F1 score with respect to the hyper parameter values used for training the model. The X-axis is Gamma value ranging from 0.001 to 5.001 and the Y-axis is C value ranging from 0.1 to 100.1.

**Figure 2:**
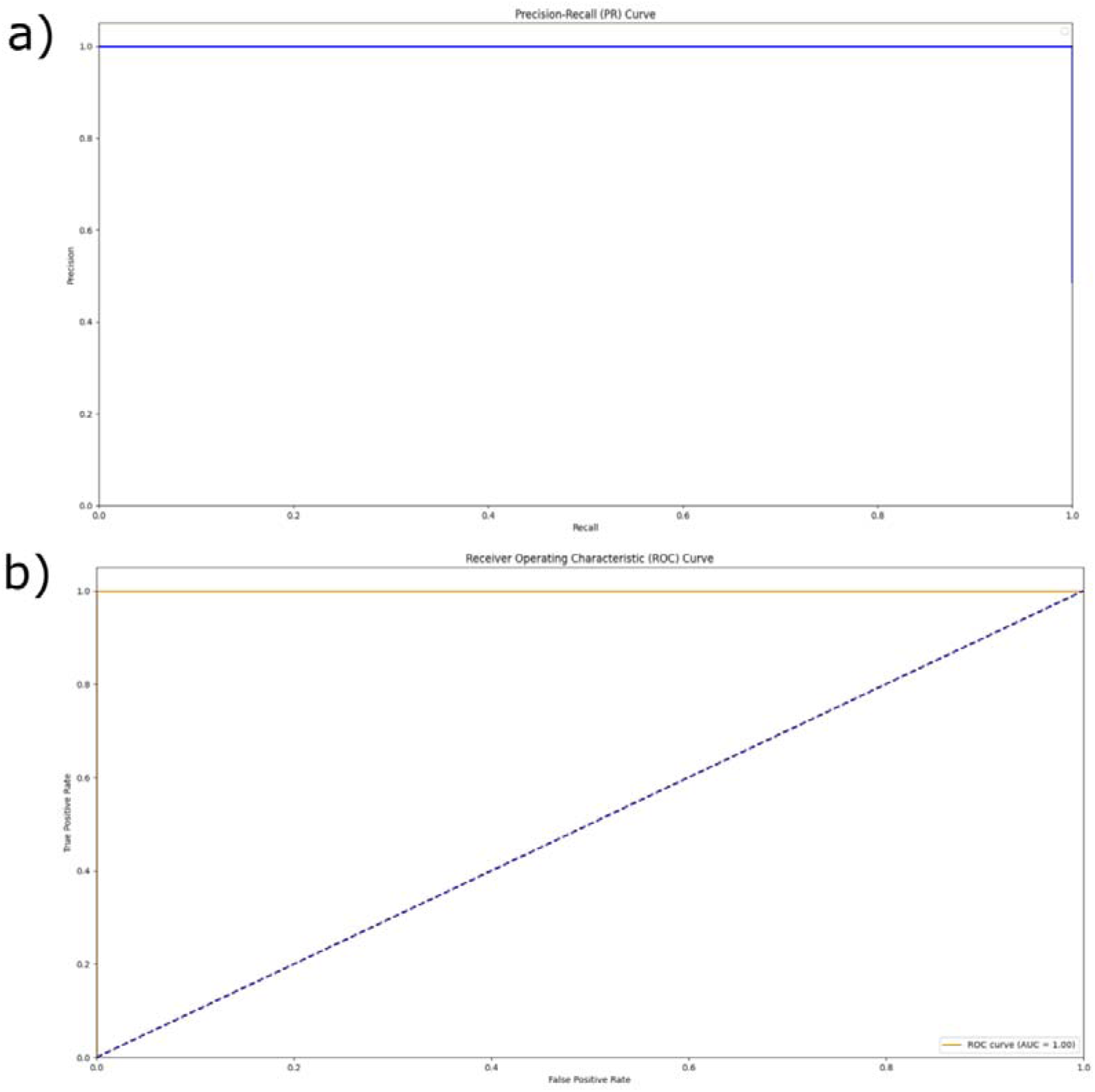
Precision-Recall curve and the Receiver Operating Characteristic (ROC) curve for the SVM model using the RBF kernel both demonstrated an AUC of 1.00. In the plots, the dotted diagonal line represents the baseline performance with an AUC of 0.5. In plot (a), the Y-axis denotes precision and the X-axis denotes recall, while in plot (b), the Y-axis represents the true positive rate and the X-axis represents the false positive rate.

### Multi-layer perceptron

We performed classification using a Multi-Layer Perceptron (MLP) model with four activation functions ReLU, tanh, identity, and logistic in combination with three solvers: adam, lbfgs, and sgd. Among the various MLP configurations, the combination of ReLU activation and adam solver delivered the best performance, achieving a maximum F1 score of 0.99, supported by a confusion matrix of 40 true positives (TP), 0 false positives (FP), 1 false negative (FN), and 37 true negatives (TN). For the identity activation function, the adam solver attained an F1 score of 0.77 on the validation set (Supplementary Figure 9) and 0.54 on the independent set (Supplementary Table 3), while the lbfgs and sgd solvers produced validation scores of 0.69 and 0.77 (Supplementary Figures 10 and 11) and dropped to 0.51 and 0.54 on the independent set, respectively. With the logistic activation, the adam solver reached an F1 score of 0.99 on the validation set (Supplementary Figure 12), but fell to 0.69 in the independent evaluation. Similarly, the lbfgs solver scored 1.00 on the validation set (Supplementary Figure 13) and 0.68 on the independent set, while the sgd variant recorded 0.71 and 0.63, respectively (Supplementary Figure 14). The ReLU activation function consistently performed well across solvers, with the adam, lbfgs, and sgd solvers achieving validation F1 scores of 0.99 (Figure 3), 1.00 (Supplementary Figure 15), and 0.97 (Supplementary Figure 16), and independent set scores of 0.76, 0.69, and 0.57, respectively (Table 2). The tanh activation function also showed strong results, where the adam, lbfgs, and sgd solvers recorded F1 scores of 0.97, 1.00, and 0.77 in the validation set (Supplementary Figures 17–19) and 0.78, 0.64, and 0.57 on the independent set (Table 2). Overall, the ReLU and tanh activation functions outperformed the identity and logistic variants across both datasets. The best-performing model achieved a precision of 0.98, recall of 0.97, and an AUC of 1.00 (Figure 4), with a confusion matrix of 40 TP, 0 FP, 2 FN, and 36 TN on the validation set, and 1304 TP, 168 FP, 257 FN, and 260 TN on the independent set.

**Figure 3:**
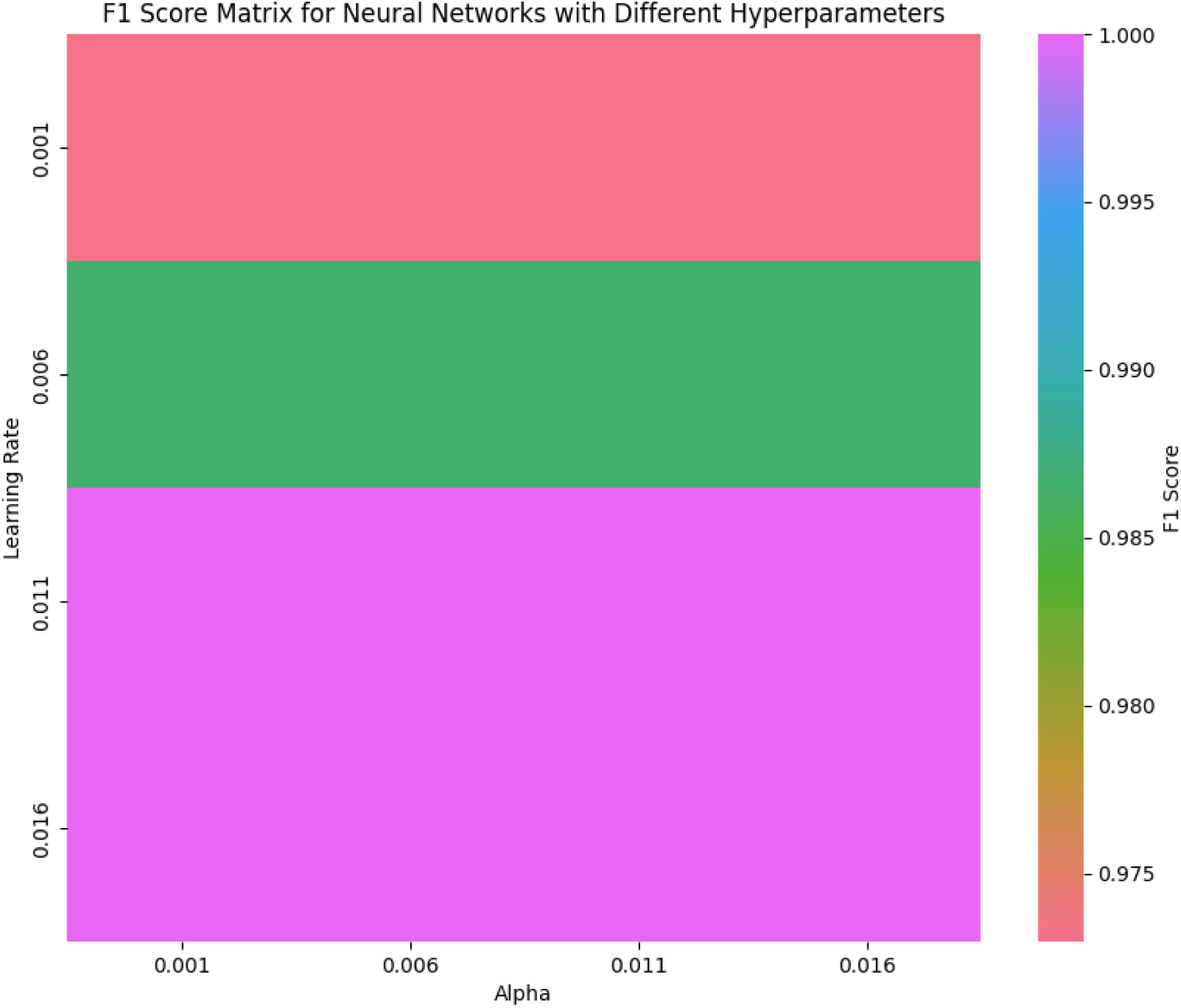
Heatmap plot of F1 score for Adam solver and Relu activation functions used in the MLP for finding the parameter values for best model. The top F1 scores obtained are as follows: Identity-adam - 0.77, Logistic-adam - 0.99, ReLU-adam - 0.99, and Tanh-adam - 0.97. The colors indicate the F1 score with respect to the hyper parameter values used for training the model. The X-axis is Alpha value ranging from 0.001 to 0.016 and the Y-axis is learning rate ranging from 0.001 to 0.016.

**Figure 4:**
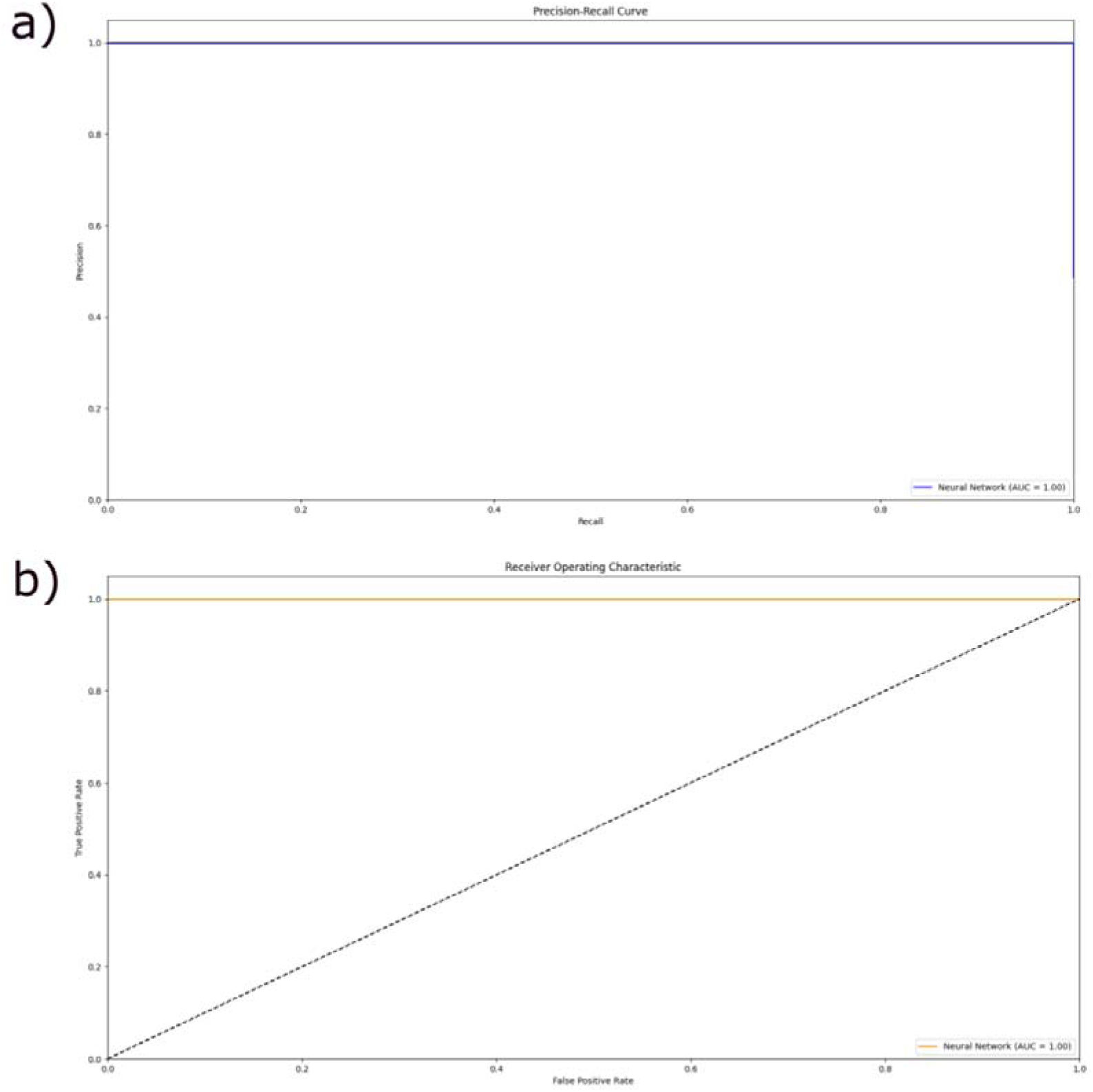
Precision-Recall curve and Receiver operating characteristic (ROC) curve with an AUC of 1.00 for Multi-layer perceptron. The dotted line represents the diagonal and AUC 0.5 of the plot. In plot (a), the Y-axis denotes precision and the X-axis denotes recall, while in plot (b), the Y-axis represents the true positive rate and the X-axis represents the false positive rate.

We evaluated the ML models generated using various algorithms, with sequence data from the CAZy families modeled using AlphaFold2. A total of 739, 3459, 34, 43, 354, 59, and 6 models were generated for AA9, AA10, AA11, AA13, AA15, AA16, and AA17, respectively. A study by Yu, X., et al. (2023) on the AA9 family of LPMOs showed chitinolytic activities. We validated the trained model with sequences from these families, dividing them into chitinolytic and cellulolytic LPMOs. The performance of the models is shown in Table 2 Supplementary table 2, and Supplementary table 3 where positive represents chitinolytic LPMOs and negative represents cellulolytic LPMOs.

In this study, we systematically evaluated multiple machine learning algorithms including Logistic Regression, Gaussian Naïve Bayes, Random Forest, Support Vector Machine (SVM), and Multi-Layer Perceptron (MLP) to classify chitinolytic and cellulolytic LPMOs using AlphaFold2-generated sequence models from various CAZy families. Among these, SVM with a radial basis function (RBF) kernel and the MLP model with ReLU and tanh activation functions demonstrated superior performance in both validation and independent datasets. SVM-RBF emerged as a robust model with an F1 score of 0.99 on the validation set and 0.74 on the independent set, coupled with an AUC of 1.00, highlighting its ability to generalize well. Similarly, MLP models using ReLU–adam and tanh–adam combinations achieved near-perfect validation metrics, including F1 scores up to 0.99 and AUC of 1.00, while maintaining moderate performance on independent datasets (F1 scores of 0.76 and 0.78, respectively). These configurations consistently outperformed others across both datasets. In contrast, although Random Forest and Gaussian Naïve Bayes delivered strong performance on the validation set (F1 score = 0.96), they failed to generalize, exhibiting extremely poor performance on the independent set due to overfitting. Logistic Regression offered modest and stable results, with liblinear and lbfgs solvers reaching the highest F1 score of 0.65 on the independent set. Overall, the results underscore the importance of evaluating models on independent datasets and tuning hyperparameters effectively. Models built using deep learning approaches like MLP and kernel-based methods such as SVM, when appropriately configured, show strong potential for distinguishing between chitinolytic and cellulolytic LPMOs. The findings advocate for the ReLU–adam MLP and SVM-RBF models as the most reliable candidates for this classification task, offering high precision, recall, and generalizability.

Upon comparing all 24 models, the SVM and Multi layer perceptron models exhibited the highest overall performance on the independent dataset. Both models showed lower false positive predictions, validating the concept that 3D topological and structural features can effectively train a model to classify protein functions.

One of the main limitations of these models lies in their inability to effectively capture the complexity of proteins that exhibit multiple enzymatic activities. When a single protein performs more than one biological function, it becomes challenging for the models to distinguish between activity-specific patterns, leading to reduced prediction accuracy. Additionally, the models were trained using a relatively limited set of descriptors, which may not comprehensively represent the structural or functional diversity of the protein sequences. This constraint can restrict the model’s ability to generalize to new, unseen data, ultimately affecting its robustness and applicability in real-world scenarios.

Using protein structural features for annotation is a novel approach that provides unique insights into the three-dimensional structure and function of proteins. As more sequences and structures are identified, future improvements in model performance are anticipated.

## Materials and Methods

We started by collecting LPMO sequence data from the CAZy database. As of January 2023, statistical data of each family on the CAZy database has been recorded (Table 3). We segregated the protein structure of LPMOs as chitinolytic and cellulolytic. We used scripts for filtering sequences by referring to other databases to clean the datasets. We used AlphaFold2 to model the amino acid sequence present in CAZy. We used in-house Python scripts to convert PDB files into STL files using PyMOL. After the conversion of PDB files to STL format we used MSMS a tool to extract structural features of a protein, and Python scripts using modules like Open3D, trimesh and NumPy 3D structural features were extracted. All the extracted features were numerical in nature and these numerical were stored in a CSV file. The features extracted are reduced surface, solvent-excluded surface, concavity, convexity, and other features describing the topological features of a 3D model. The CSV file containing extracted features was used to find the significant feature that can be used to train a model, using Ensemble feature selection (EFS) an R programming-based package which helps in finding significant features by performing a series of eight different classifications to each descriptor to find a normalized score between the descriptors. This normalized score ranges from 0 to 1 and is known as the ensemble score. A score of more than 0.5 in ensemble score would be a significant feature that can be used to train a model. After getting the significant feature, we developed machine learning-based models which employ more than one approach to categorize a given set of structural data of a protein.

**Table 3:** Number of sequence and structural information available in the CAZy database.

| AA families | Number of sequence information in CAZY.org | Number of structural information in CAZY.org |
| --- | --- | --- |
| AA9 | 1336 | 101 |
| AA10 | 11503 | 51 |
| AA11 | 367 | 3 |
| AA13 | 63 | 8 |
| AA14 | 136 | 1 |
| AA15 | 498 | 1 |
| AA16 | 103 | 1 |
| AA17 | 93 | 1 |
| **Total** | **14099** | **167** |

We employed support vector machine (SVM), random forest, logistic regression, Naïve Bayes and multi layer perceptron, predictions after calculating the 3D model features. The models were cross-validated with validation and test set and the models were optimized with the help of F1 score. Finally, the models generated were evaluated by performance metrics such as precision, recall, and accuracy.

### PyMOL

PyMOL is a software used mainly for visualization and analyzing protein structures (Schrödinger, L., & DeLano, W., 2020). It can be used with either a graphical user interface or it can also be used via a command line. Python-based scripting features are an in-built feature in PyMOL which helps users in automating tasks, creating workflows, and even implementing PyMOL as a part of a pipeline. The commercial version of PyMOL is required for the conversion of the PDB protein file into a 3D model file.

### Python

Python is one of the elegant and widely used programming languages. Python is known for its ability to simplify and optimize the performance of a script. Python offers various libraries and frameworks that enhance its capabilities for specific tasks. Python helps in automating a series of processes and creating a workflow by integrating different modules, and shell processes. We used various Python modules in developed Python scripts for performing data analysis, generating 3D models, extracting features, generating Machine learning models and evaluating ML models. We used Python scripts to run on PyMOL to create 3D models. For the data analysis, visualization, and pre-processing modules like NumPy, Pandas and Matplotlib were used. Extraction of 3D features was done by using modules like open3D (Zhou, Q. *et al*. 2018), trimesh, and numpy-stl. For the preparation of ML models python module scikit-learn was used (Fabian Pedregosa *et al*. 2011).

### trimesh, numpy-stl and open3D

These are Python modules used to retrieve features from a 3D structure, and the features extracted provide valuable insights into the geometry and physical properties of the model. The geometric and structural properties of 3D models can be characterized using several important features. Scale refers to the ability of a 3D object to uniformly adjust its size along the X, Y, and Z axes while maintaining its original shape and proportions. This is essential for comparing models of varying dimensions. The surface area represents the total area covered by the outer surface of the 3D model, while the mesh volume indicates the overall volume enclosed by the model’s surface. Complementing this, the convex volume measures the volume of the smallest convex shape that can fully contain the model. The relationship between the model’s surface features is captured through concavity and convexity where concavity is the ratio of the concave surface area to the total surface area, and convexity is the ratio of the convex surface area to the total. Compactness is a metric that reflects how tightly packed or dense the 3D structure is, offering insights into the structural integrity of the model. Additionally, sphericity and asphericity describe the shape characteristics of the model’s surface sphericity being the percentage of the surface that is spherical in nature, and asphericity representing the portion that is more linear or deviates from spherical form. Together, these features provide a comprehensive understanding of the physical and spatial attributes of 3D objects.

### MSMS

MSMS (Molecular Surface, Molecular Surface Package) is a software package that provides an efficient and accurate method for computing molecular surfaces in 3D space. The "Reduced surface" algorithm is a specific feature within MSMS that offers an efficient approach for calculating the molecular surface of a biomolecule. The combination of MSMS’s "Reduced surface" algorithm and its comprehensive set of features makes it a valuable tool for extracting features from biomolecules (Sanner *et al* 1996).

Both positive and negative datapoint was separately given as input in a Python script developed in-house which uses previously mentioned Python modules and MSMS to generate 34 distinct descriptors.

### Reduced surface

In terms of MSMS (Michel Sanner’s Molecular Surface), the reduced surface refers to a method by which the complex 3D structure of a protein is simplified and optimized to allow for more efficient and meaningful extraction of structural features. This reduction enables faster computations and more accurate analysis of the protein’s surface geometry. The following features represent numerical, quantitative descriptors derived from such a reduced 3D model, faces: This refers to the number of polygonal surfaces (usually triangles) that make up the outer surface of the structure after it has been processed. Edges: Represents the total number of line segments that form the boundaries between faces in the optimized structure. Free Edges: These are the edges that are not shared between two faces essentially, they lie on the boundary of holes or incomplete regions of the mesh. Vertices: Denotes the number of distinct corner points where edges meet in the 3D model. Genus: A topological property that characterizes the surface by counting the number of holes or handles it possesses (e.g., a sphere has a genus of 0, a torus has a genus of 1). These descriptors provide critical insights into the topology and geometry of the protein structure, serving as the foundation for downstream structural and functional analyses.

### Analytical solvent excluded surface

The Analytical Solvent Excluded Surface (ASES) is a sophisticated computational method designed to calculate the molecular surface area that is inaccessible to solvent molecules. Unlike simpler surface estimation techniques, ASES simulates the interaction between a molecular structure and a solvent probe, producing a highly accurate and detailed depiction of the molecule’s true physical boundaries. This level of detail is essential for understanding how proteins interact with other molecules, including ligands and solvents, particularly in the context of drug design or enzymatic function. In this representation, several structural and surface-related features are extracted from the model. The number of faces refers to the polygonal divisions across the solvent-excluded surface, while edges and vertices describe the mesh structure that defines how these faces connect in three-dimensional space. The surface area of the edges (S_edge) and the surface area of the vertices (S_vertex) provide quantitative insights into the geometric contribution of these components to the overall structure. Moreover, the radius of the solvent probe (R_H) used in the simulation plays a crucial role in defining which regions of the molecule are considered inaccessible, while the probe density (C_H) affects the resolution and precision of the surface calculation. Finally, the genus a topological measure captures the complexity of the surface by indicating the number of holes or handles, contributing to a deeper understanding of molecular topology. Altogether, ASES-derived features help generate a nuanced, quantitative map of molecular surfaces that is invaluable for structural biology and bioinformatics analyses.

### Analytical surface area

The analytical surface area in MSMS (Michel Sanner’s Molecular Surface) represents a precise method for calculating the total surface area of a molecule using analytical techniques. This approach involves computing the surface area contributed by each individual atom and then summing them to derive the molecule’s overall surface characteristics. Such detailed surface calculations are crucial in molecular modeling, particularly for understanding interactions with solvents, ligands, and other biomolecules. Several structural features are derived through this method. The Reent region identifies concave pockets within the molecular surface, which are often critical in binding interactions. The Toric feature captures toroidal or doughnut-like formations within the molecule, indicative of complex structural topologies. The Contact feature highlights areas where atoms or molecules are in close proximity, reflecting potential interaction sites. Two important quantitative measures derived from this method are the SES (Solvent Excluded Surface area), which represents the surface inaccessible to solvent molecules, and the SAS (Solvent Accessible Surface area), denoting the area accessible to solvent probes. Together, these features provide a comprehensive and accurate description of a molecule’s surface topology, essential for structural and functional protein analysis.

### Triangulation

In MSMS (Molecular Surface and Volume), triangulation refers to the computational process of constructing a mesh representation of a molecular surface by dividing it into a series of interconnected triangles. This technique is employed to approximate the surface’s geometric and topological features with a high level of precision, facilitating efficient visualization, analysis, and further computational modeling. The triangulated surface is characterized by several quantitative attributes. The vertices represent the distinct corner points of each triangle on the mesh, while the edges are the lines connecting these vertices. The faces are the triangular patches that collectively form the surface of the molecular model. A topological descriptor, known as T_genus, captures the intrinsic properties of the surface such as the number of holes or handles helping to understand the complexity of the molecular shape. Lastly, density in this context measures how tightly the features or geometric properties are distributed across the surface, indicating the resolution and compactness of the triangulated mesh. This level of detail is essential for accurate structural analyses and simulations in molecular biology and computational chemistry.

### Numerical volumes and area

In MSMS, the solvent-excluded surface (SES) is determined using the previously mentioned analytical methods to model the regions of a molecular structure that are inaccessible to solvent molecules. This process generates a more accurate representation of the molecule’s true boundary by accounting for the spatial constraints imposed by solvent interaction. The computed SES provides valuable insights into molecular stability, interaction sites, and structural compactness. Two important numerical features derived from this analysis are the SES volume and the SES area. The SES volume quantifies the three-dimensional space enclosed by the solvent-excluded surface, essentially representing the molecular core that cannot be penetrated by solvent molecules. Meanwhile, the SES area calculates the total external surface area of this region, offering a precise measure of how much of the molecular surface is shielded from solvent interaction. Together, these features serve as critical indicators in structural bioinformatics, drug design, and protein-ligand interaction studies.

### Model Generation

Features extracted were used as input for in-house Python scripts. These scripts implemented various machine learning methods using libraries such as **scikit-learn**, **pandas**, **matplotlib**, **seaborn**, and **NumPy**.

Logistic regression is a statistical model often used for binary classification. It estimates the probability of a binary outcome based on input features. Six solvers were tested to identify the best-fit logistic regression model for this dataset (Cox, D. R., 1958).

Naïve Bayes, a probabilistic classifier based on Bayes’ theorem, was implemented under the assumption of feature independence (Feng et al., 2013)

Random Forest, an ensemble-based algorithm combining multiple decision trees, was applied to enhance model accuracy and reduce overfitting (Chen et al., 2012).

The primary objective of SVM is to find the optimal hyperplane that separates data points in a high-dimensional feature space into distinct classes. This hyperplane maximizes the margin between the closest data points of each class, known as support vectors. Parameters such as kernel functions significantly influence the decision boundary and were tested to identify the best-fit model (Byvatov, E., & Schneider, G., 2003).

Neural networks (NN), inspired by the structure and function of the human brain, consist of interconnected artificial neurons organized into layers. NNs excel at identifying complex patterns and relationships in data, making them suitable for tasks such as classification and regression. Various solvers and activation functions were tested to determine the best configuration for this dataset (McCulloch, W. S., & Pitts, W., 1943).

### Validation of models

Model performance was assessed using **precision**, **recall**, **F1 score**, and **accuracy** metrics on validation and independent datasets.

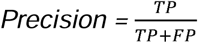

Precision measures the proportion of correctly predicted positive instances among all instances predicted as positive.

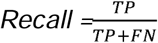

Recall measures the proportion of actual positive instances correctly predicted by the model.

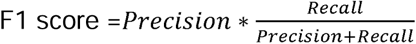

The F1 score balances precision and recall, offering a comprehensive measure of a model’s performance.

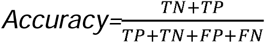

Accuracy measures the proportion of correct predictions out of the total predictions made.

## Conclusion

We have successfully implemented a Machine learning model, where 3D structural features extracted from a protein file can be used for performing annotation of protein function. As a proof of concept, this was implemented in LPMOs, where chitinolytic and cellulolytic LPMOs are major biomass sources for partial replacement of fossil fuels or petroleum products and Machine learning models were able to perform well at categorizing LPMOs. The SVM algorithm with Radial basis function (*rbf*) as a kernel was the best model obtained for this dataset with the highest accuracy of 0.98 and the Multi-layer perceptron algorithm with a solver as *adam* and activation function tanh as also performed well for the dataset with an accuracy of 0.97. Currently, LPMOs that act only on one type of substrate are used to evaluate. Thus, limiting the performance of the model when the dataset contains protein that can act on more than one substrate.

## Acknowledgements

The authors acknowledge SASTRA Deemed to be University for infrastructural support.

## Credit Author

AM did the Data Curation, Data Analysis, Modeling, Validation, and wrote the first draft of the manuscript. RMY did the Conceptualization, Supervision, Validation, and finalized the manuscript.

## Data and Code Availability

The scripts, training data, and trained models are available for use at the following Zenodo repository: Yennamalli, R. (2025). LPMO_classifier.

Zenodo. https://doi.org/10.5281/zenodo.15748573

