## Supplementary Figures for "Machine learning-based structural classification of lytic polysaccharide monooxygenases"

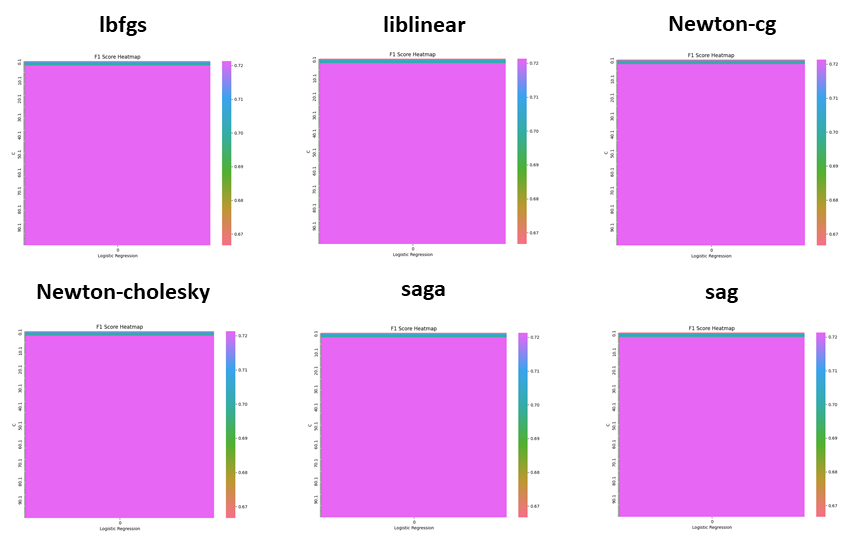

**Supplementary Figure 1.** Heatmap plot of F1 score of six solvers used in logistic regression for finding the parameter value of the best model. The optimized F1 score obtained for each solver are as follows. The lbfgs solver gave a F1 score value of 0.62, The liblinear solver gave a F1 score of 0.64, The Newton-cg solver gave a F1 score value of 0.62, The Newton-Cholesky solver gave a F1 score value of 0.62, The Saga solver gave a F1 score value of 0.32, and The Sag solver gave a F1 score value of 0.32.

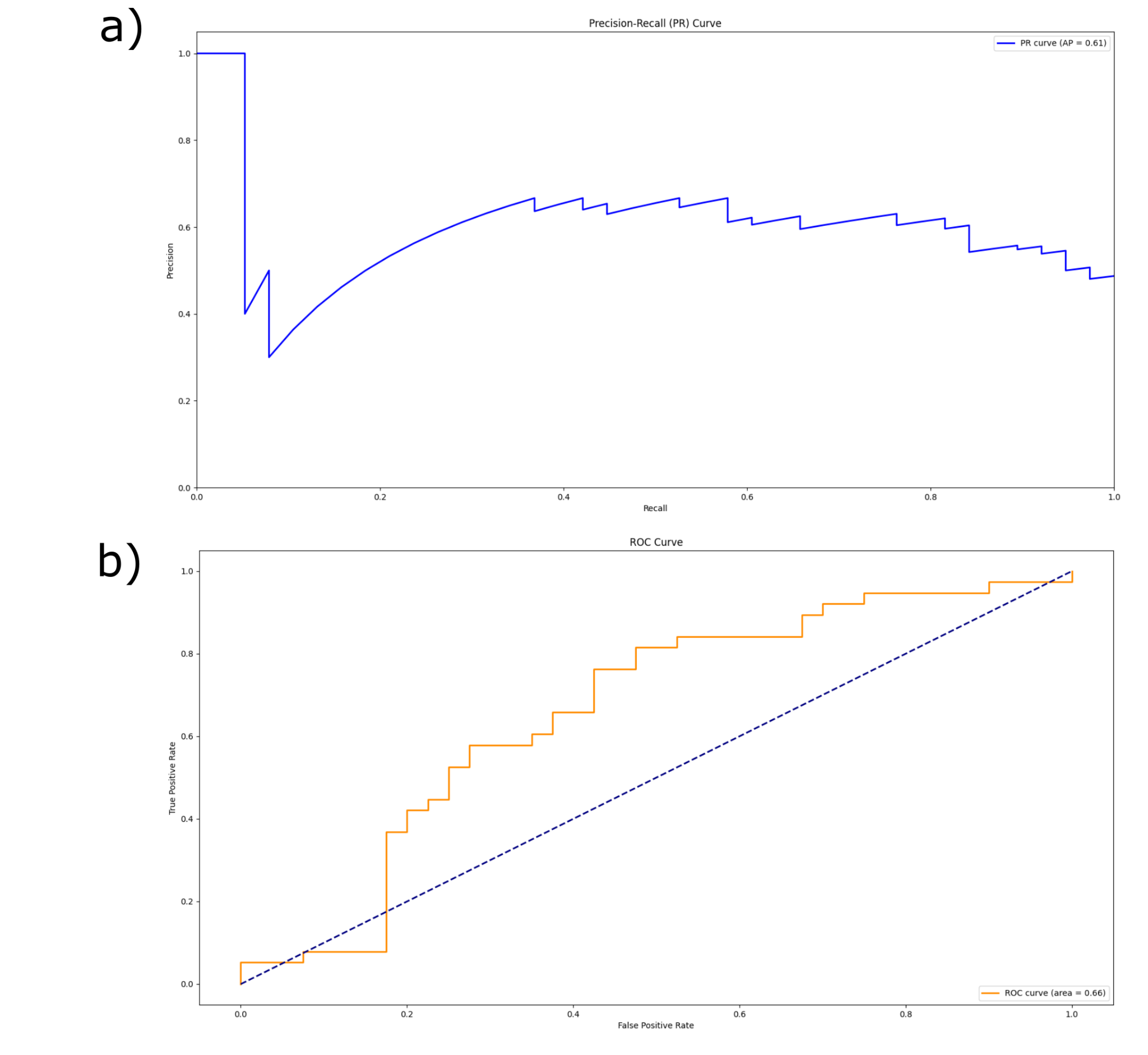

**Supplementary Figure 2. Precision-Recall curve and Receiver operating characteristic (ROC) curve with an AUC of 0.66 for logistic regression model.** The dotted line represents the diagonal and AUC 0.5 of the plot. . In plot (a), the Y-axis denotes precision and the X-axis denotes recall, while in plot (b), the Y-axis represents the true positive rate and the X-axis represents the false positive rate.

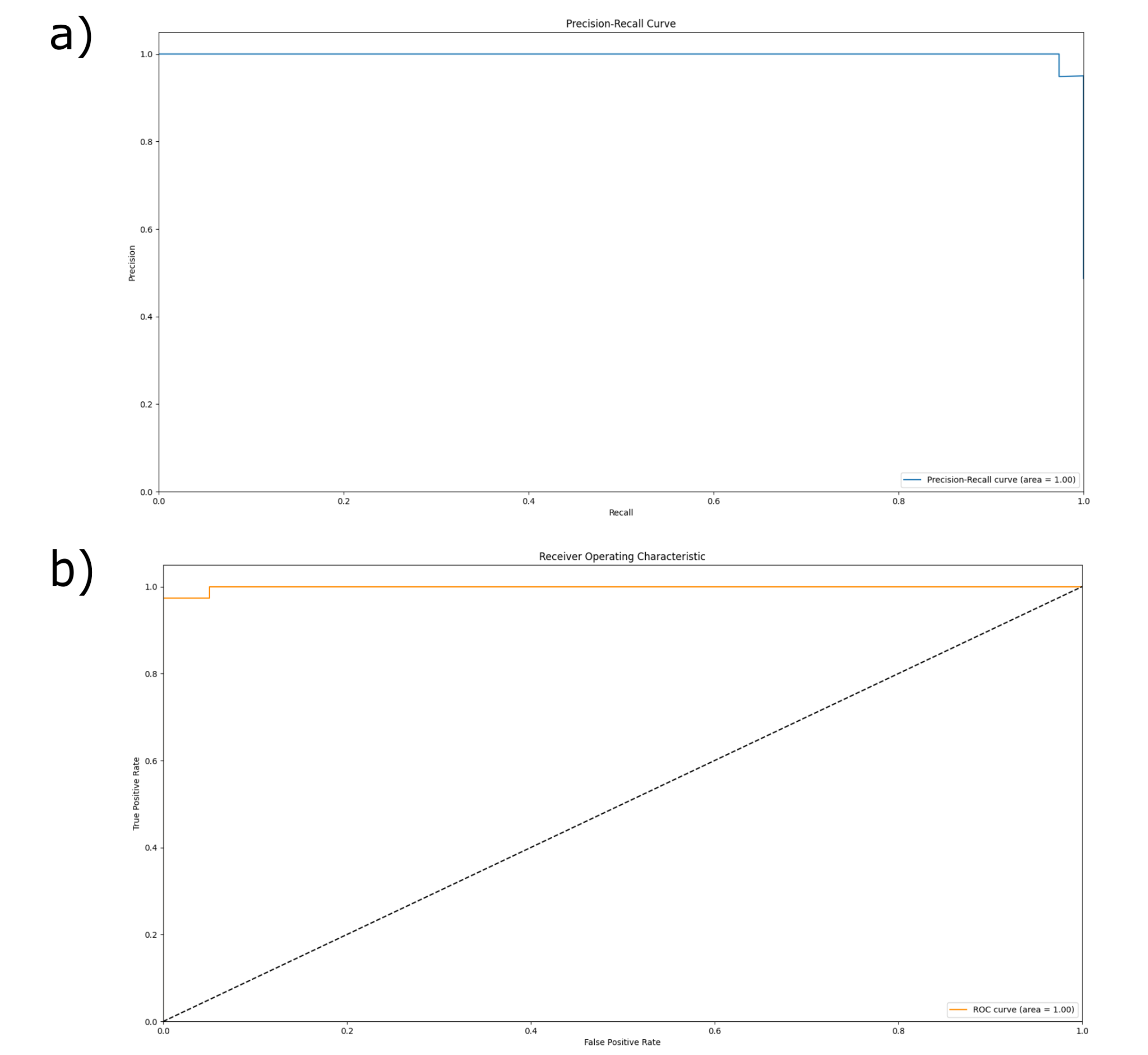

**Supplementary Figure 3. Precision-Recall curve and Receiver operating characteristic (ROC) curve with an AUC of 0.94 for the** Gaussian **Naïve Bayes model.** The dotted line represents the diagonal and AUC 0.5 of the plot. In plot (a), the Y-axis denotes precision and the X-axis denotes recall, while in plot (b), the Y-axis represents the true positive rate and the X-axis represents the false positive rate.

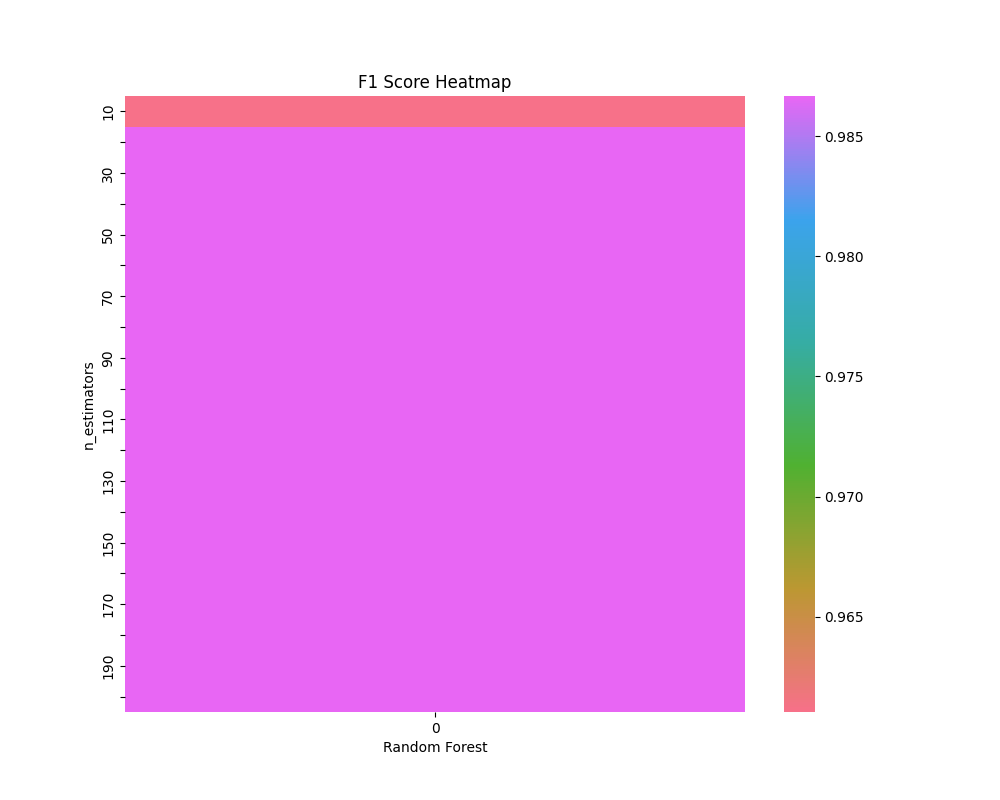

**Supplementary Figure 4. Heatmap plot of F1 score values for Random Forest with a maximum value of 0.96.** The colors indicate the F1 score with respect to the hyper parameter value used for training the model. The Y-axis is n_estimator value ranging from1-200. **a parameter and the models fit well after the value of n_estimator as 34.**

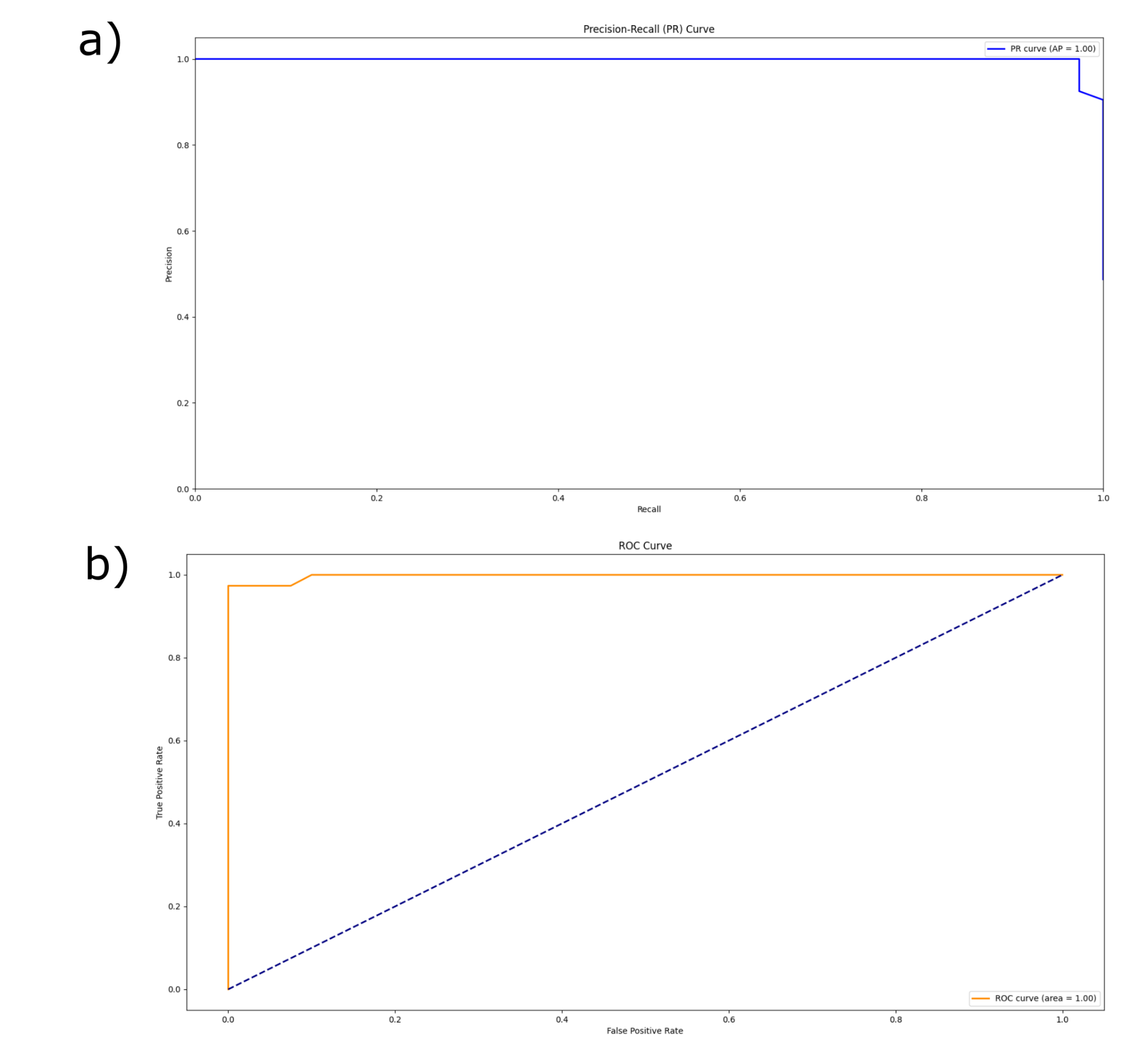

**Supplementary Figure 5.** **Precision-Recall curve and Receiver operating characteristic (ROC) curve with an AUC of 1.00 for the Random Forest model.** The dotted line represents the diagonal and AUC 0.5 of the plot. In plot (a), the Y-axis denotes precision and the X-axis denotes recall, while in plot (b), the Y-axis represents the true positive rate and the X-axis represents the false positive rate.

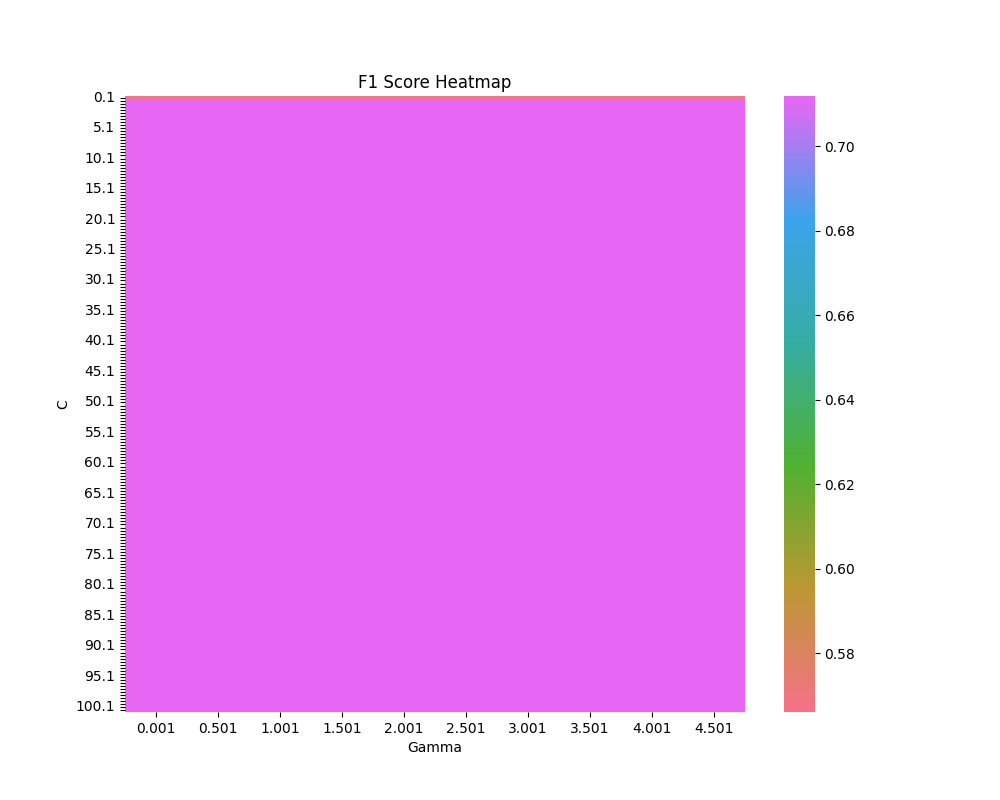

**Supplementary Figure 6:** **Heatmap plot of F1 score values for linear kernel in SVM with a maximum value of 0.77.** The colors indicate the F1 score with respect to the hyper parameter values used for training the model. The X-axis is Gamma value ranging from 0.001 to 5.001 and the Y-axis is C value ranging from 0.1 to 100.1.

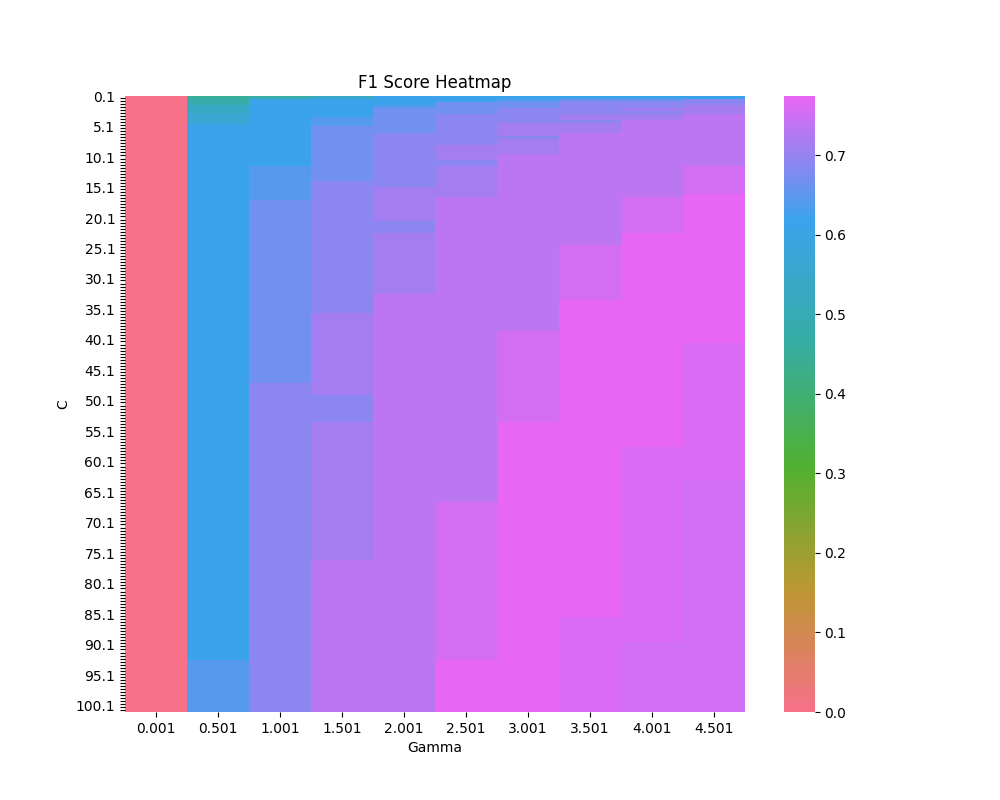

**Supplementary Figure 7:** **Heatmap plot of F1 score values for polynomial kernel in SVM with a maximum value of 0.81.** The colors indicate the F1 score with respect to the hyper parameter values used for training the model. The X-axis is Gamma value ranging from 0.001 to 5.001 and the Y-axis is C value ranging from 0.1 to 100.1.

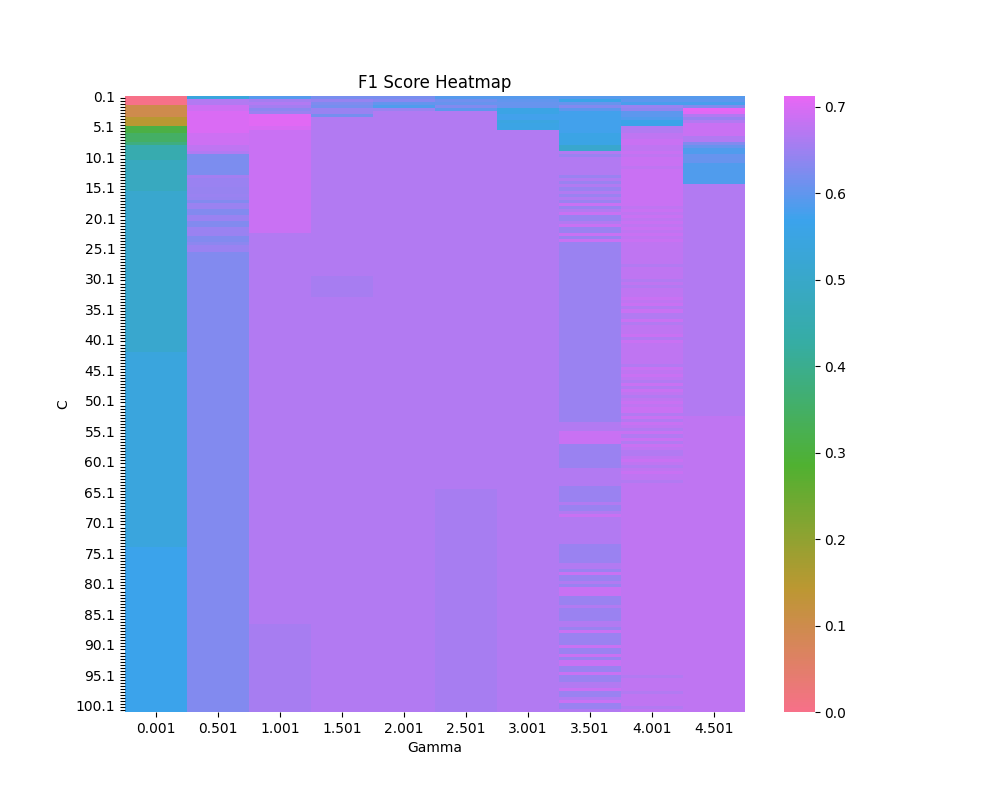

**Supplementary Figure 8:** **Heatmap plot of F1 score values for sigmoid kernel in SVM with a maximum value of 0.73.** The colors indicate the F1 score with respect to the hyper parameter values used for training the model. The X-axis is Gamma value ranging from 0.001 to 5.001 and the Y-axis is C value ranging from 0.1 to 100.1.

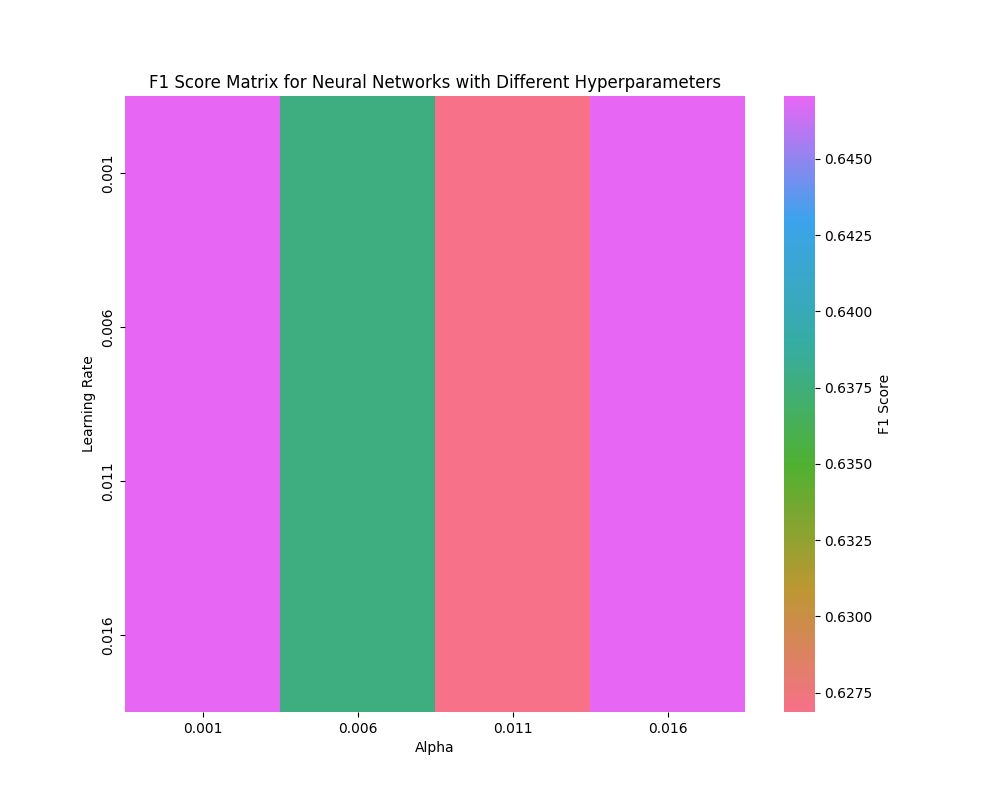

**Supplementary Figure 9:** **Heatmap plot of F1 score values for multi layer perceptron using identity activation function and adam solver with a maximum value of 0.77.** The colors indicate the F1 score with respect to the hyper parameter values used for training the model. The X-axis is Alpha value ranging from 0.001 to 0.016 and the Y-axis is learning rate ranging from 0.001 to 0.016.

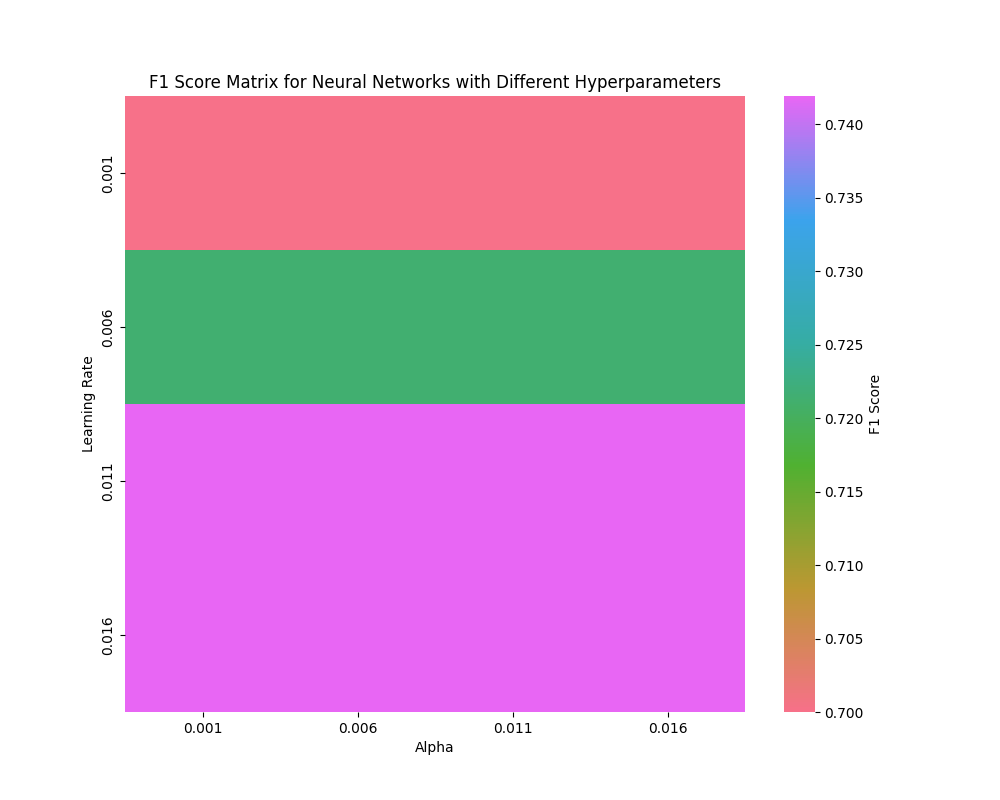

**Supplementary Figure 10:** **Heatmap plot of F1 score values for multi layer perceptron using identity activation function and lbfgs solver with a maximum value of 0.69.** The colors indicate the F1 score with respect to the hyper parameter values used for training the model. The X-axis is Alpha value ranging from 0.001 to 0.016 and the Y-axis is learning rate ranging from 0.001 to 0.016.

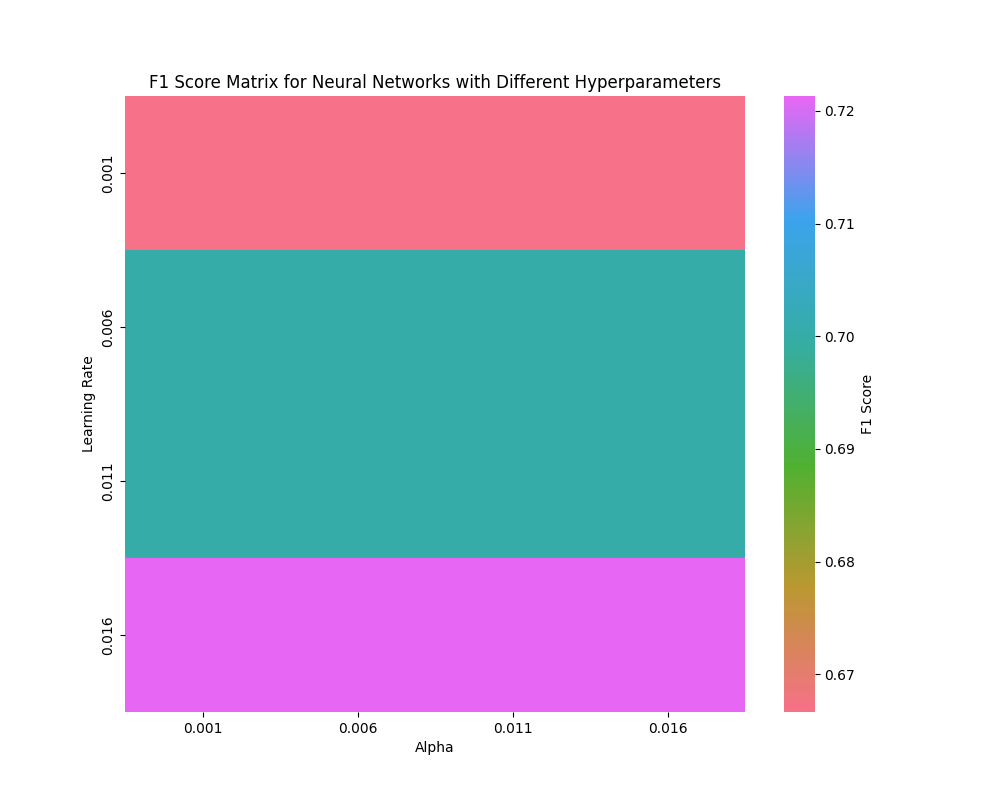

**Supplementary Figure 11:** **Heatmap plot of F1 score values for multi layer perceptron using identity activation function and sgd solver with a maximum value of 0.77.** The colors indicate the F1 score with respect to the hyper parameter values used for training the model. The X-axis is Alpha value ranging from 0.001 to 0.016 and the Y-axis is learning rate ranging from 0.001 to 0.016.

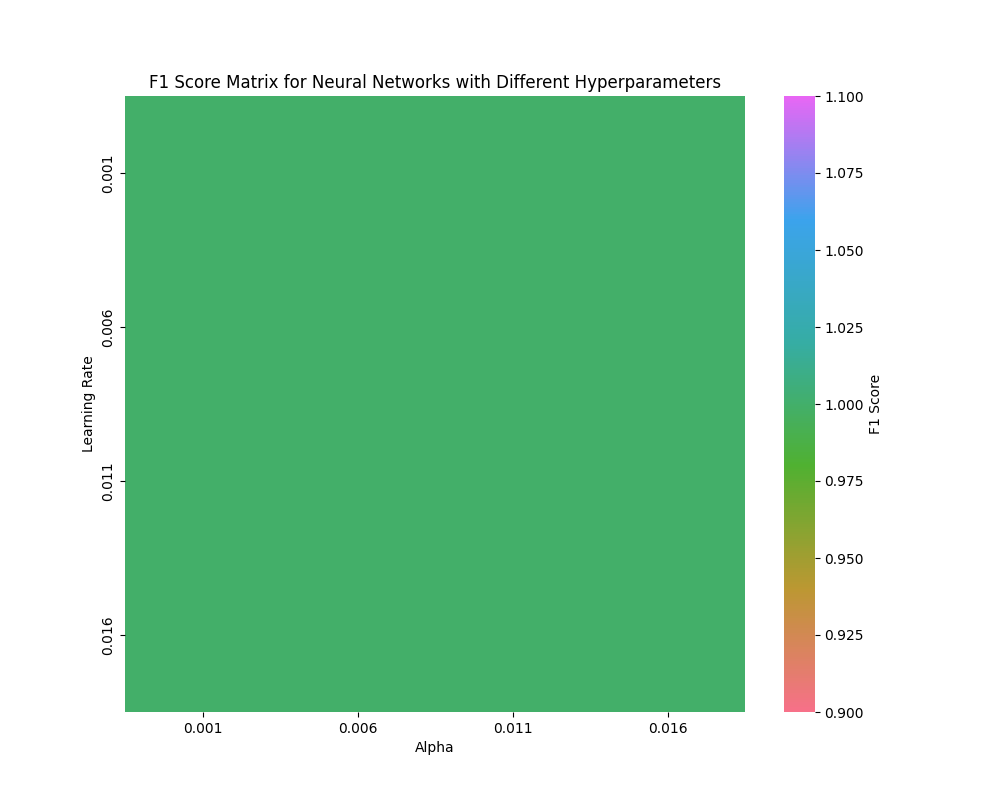

**Supplementary Figure 12:** **Heatmap plot of F1 score values for multi layer perceptron using logistic activation function and adam solver with a maximum value of 0.99.** The colors indicate the F1 score with respect to the hyper parameter values used for training the model. The X-axis is Alpha value ranging from 0.001 to 0.016 and the Y-axis is learning rate ranging from 0.001 to 0.016.

h
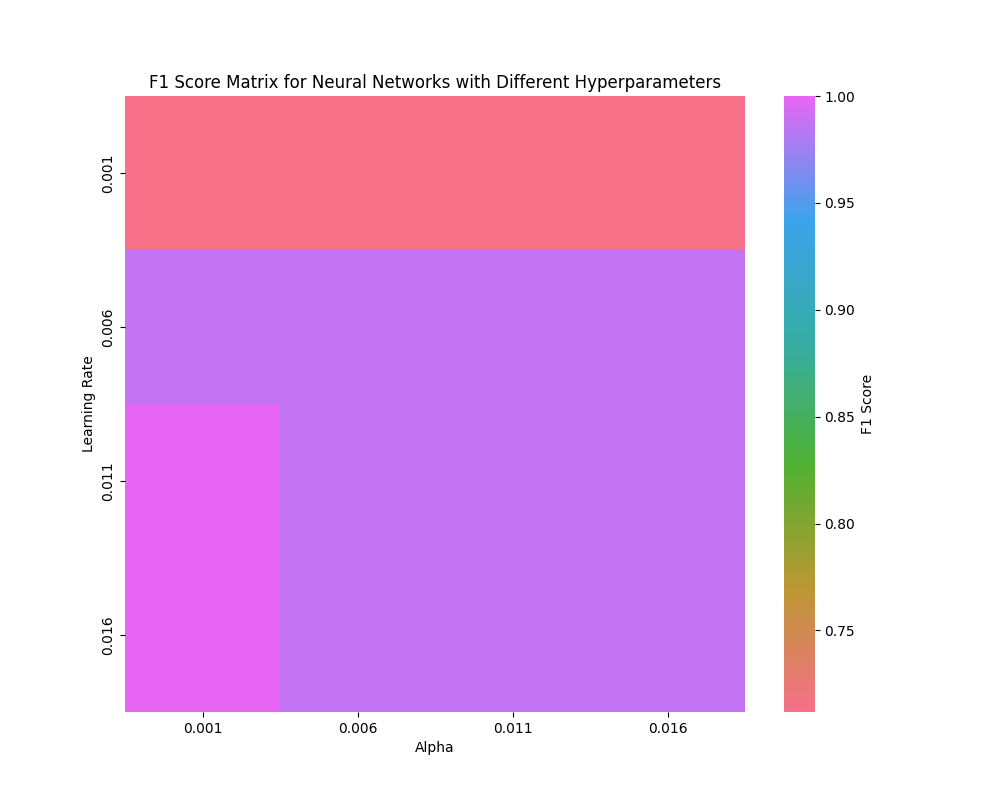

**Supplementary Figure 13:** **Heatmap plot of F1 score values for multi layer perceptron using logistic activation function and lbfgs solver with a maximum value of 1.00.** The colors indicate the F1 score with respect to the hyper parameter values used for training the model. The X-axis is Alpha value ranging from 0.001 to 0.016 and the Y-axis is learning rate ranging from 0.001 to 0.016.

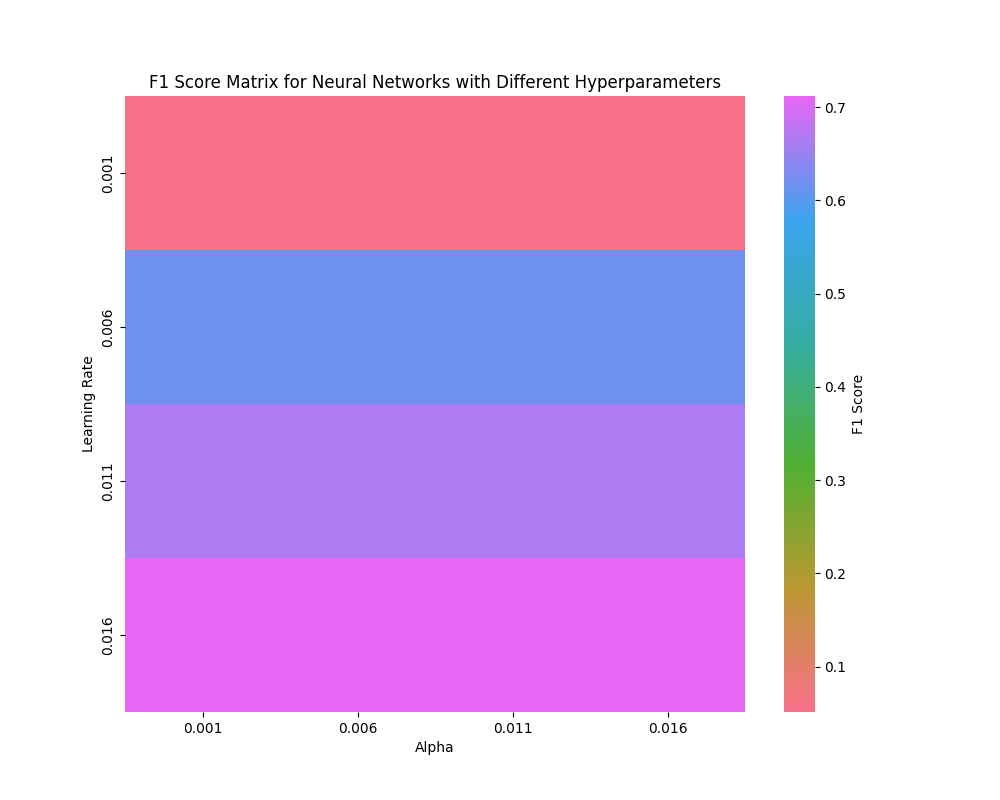

**Supplementary Figure 14:** **Heatmap plot of F1 score values for multi layer perceptron using logistic activation function and sgd solver with a maximum value of 0.71.** The colors indicate the F1 score with respect to the hyper parameter values used for training the model. The X-axis is Alpha value ranging from 0.001 to 0.016 and the Y-axis is learning rate ranging from 0.001 to 0.016.

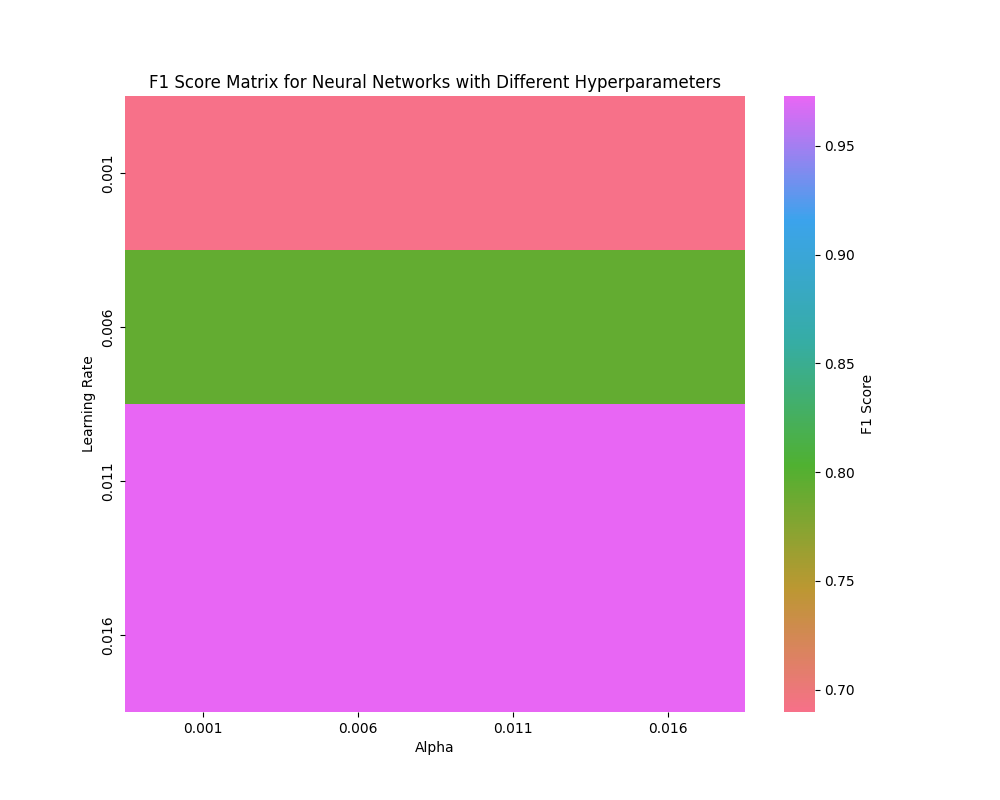

**Supplementary Figure 15:** **Heatmap plot of F1 score values for multi layer perceptron using relu activation function and lbfgs solver with a maximum value of 1.00.** The colors indicate the F1 score with respect to the hyper parameter values used for training the model. The X-axis is Alpha value ranging from 0.001 to 0.016 and the Y-axis is learning rate ranging from 0.001 to 0.016.

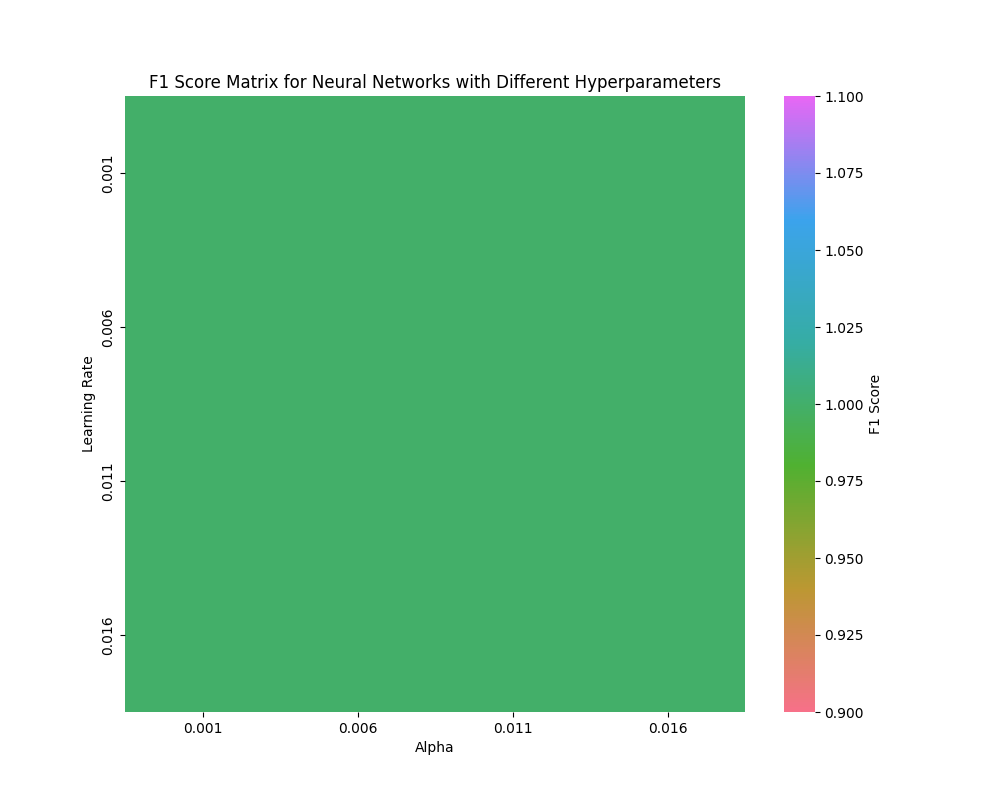

**Supplementary Figure 16:** **Heatmap plot of F1 score values for multi layer perceptron using relu activation function and sgd solver with a maximum value of 0.97.** The colors indicate the F1 score with respect to the hyper parameter values used for training the model. The X-axis is Alpha value ranging from 0.001 to 0.016 and the Y-axis is learning rate ranging from 0.001 to 0.016.

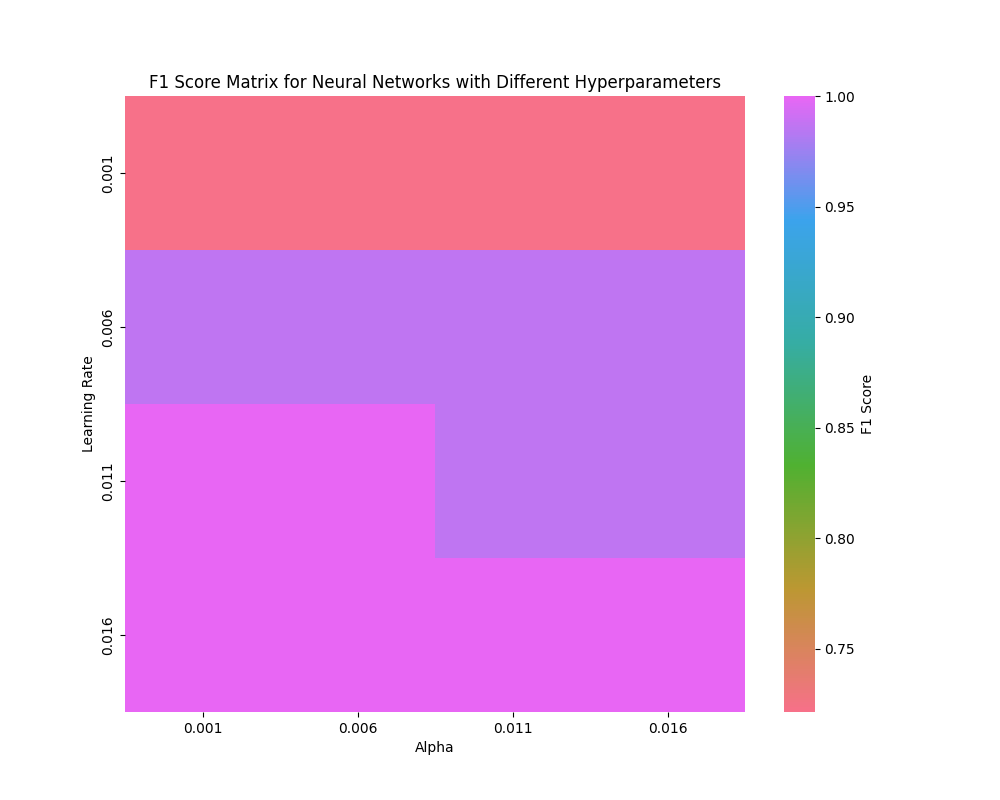

**Supplementary Figure 17:** **Heatmap plot of F1 score values for multi layer perceptron using tanh activation function and adam solver with a maximum value of 0.97.** The colors indicate the F1 score with respect to the hyper parameter values used for training the model. The X-axis is Alpha value ranging from 0.001 to 0.016 and the Y-axis is learning rate ranging from 0.001 to 0.016.

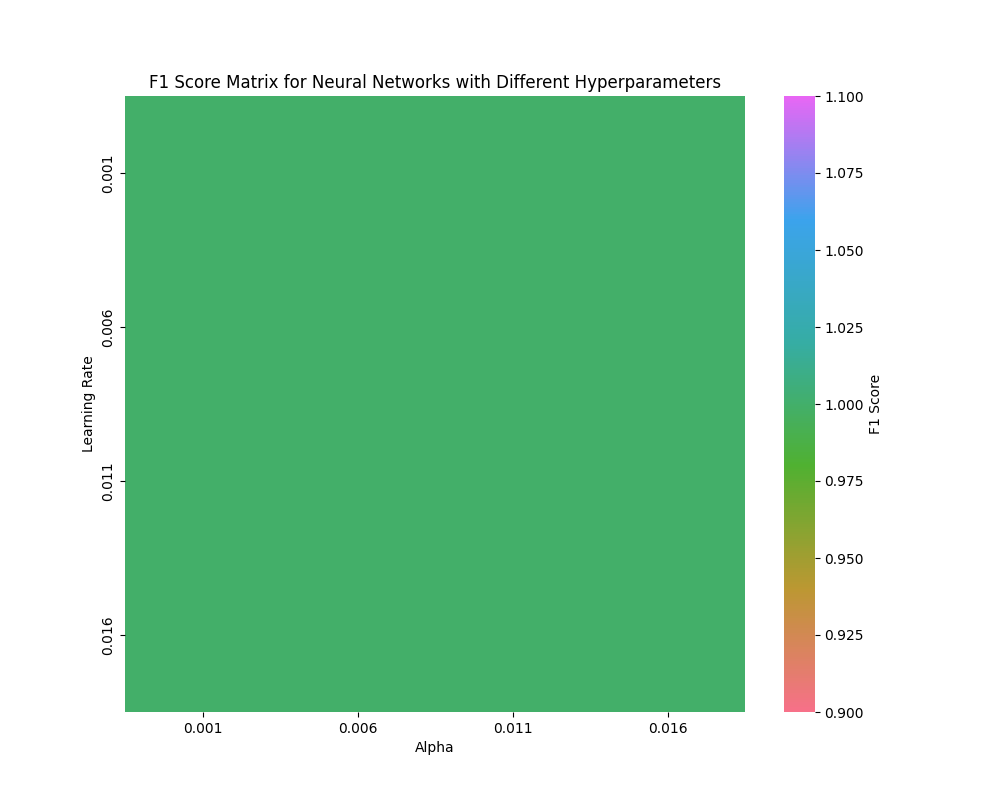

**Supplementary Figure 18:** **Heatmap plot of F1 score values for multi layer perceptron using tanh activation function and lbfgs solver with a maximum value of 1.00.** The colors indicate the F1 score with respect to the hyper parameter values used for training the model. The X-axis is Alpha value ranging from 0.001 to 0.016 and the Y-axis is learning rate ranging from 0.001 to 0.016.

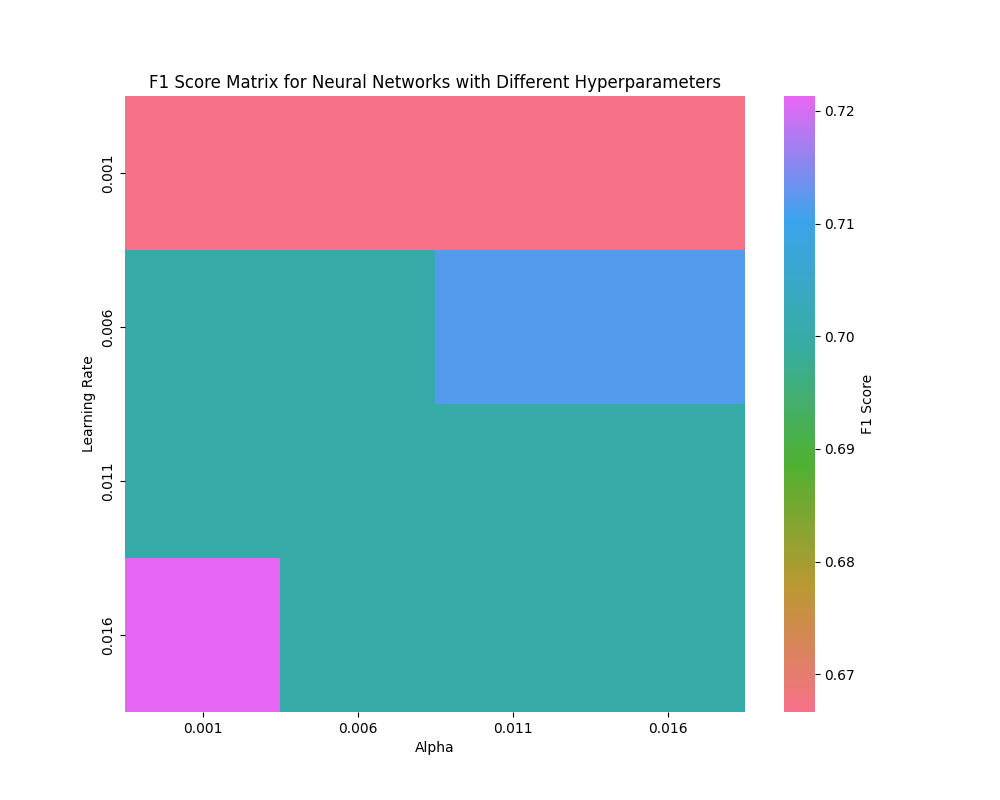

**Supplementary Figure 19:** **Heatmap plot of F1 score values for multi layer perceptron using tanh activation function and sgd solver with a maximum value of 0.77.** The colors indicate the F1 score with respect to the hyper parameter values used for training the model. The X-axis is Alpha value ranging from 0.001 to 0.016 and the Y-axis is learning rate ranging from 0.001 to 0.016.

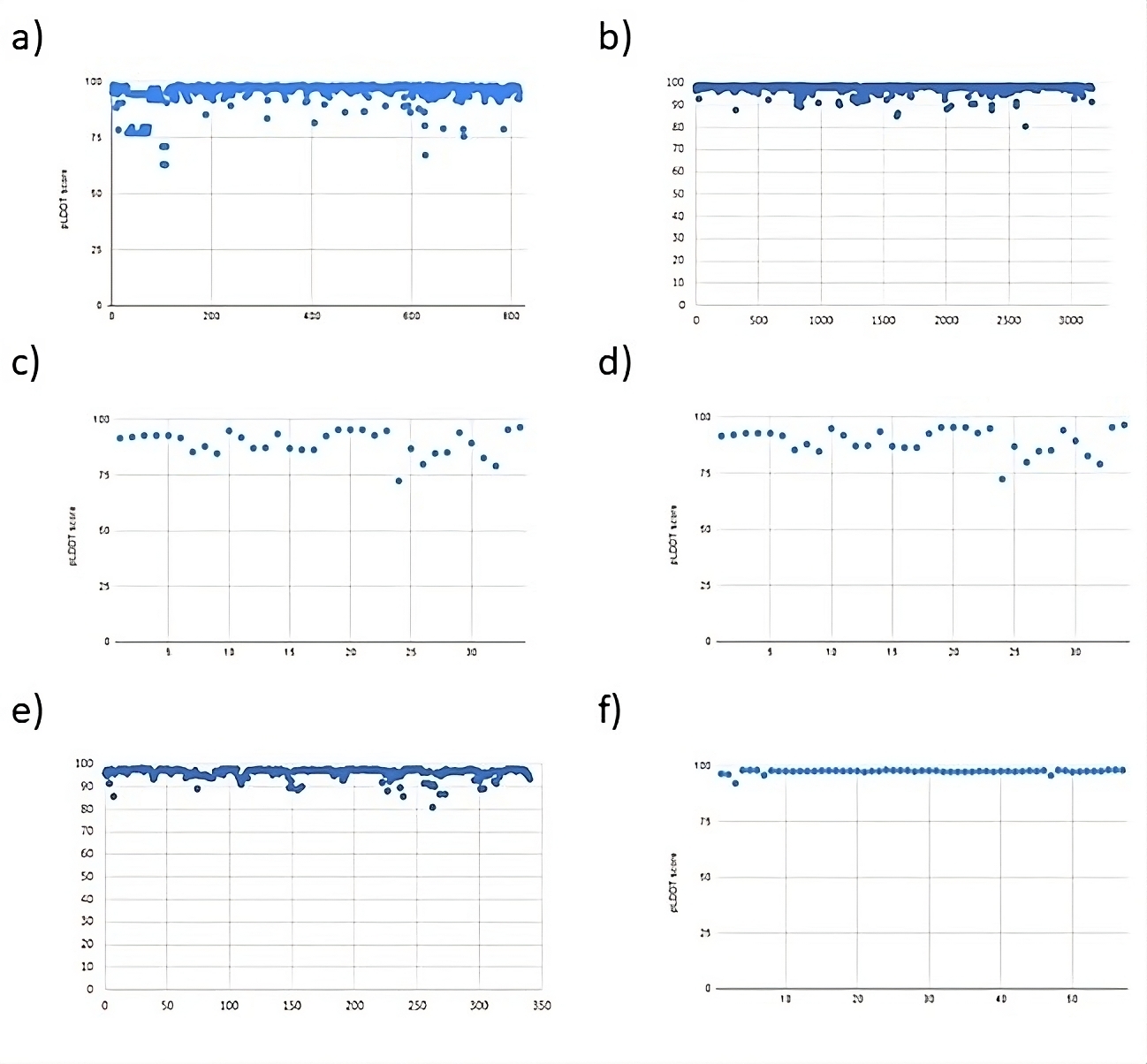

**Supplementary Figure 20. A cluster graph representing the pLDDT score of the predicted protein structures of the LPMO family. a representing AA9, b representing AA10, c representing AA11, d representing AA13, e representing AA15 and f representing AA16**

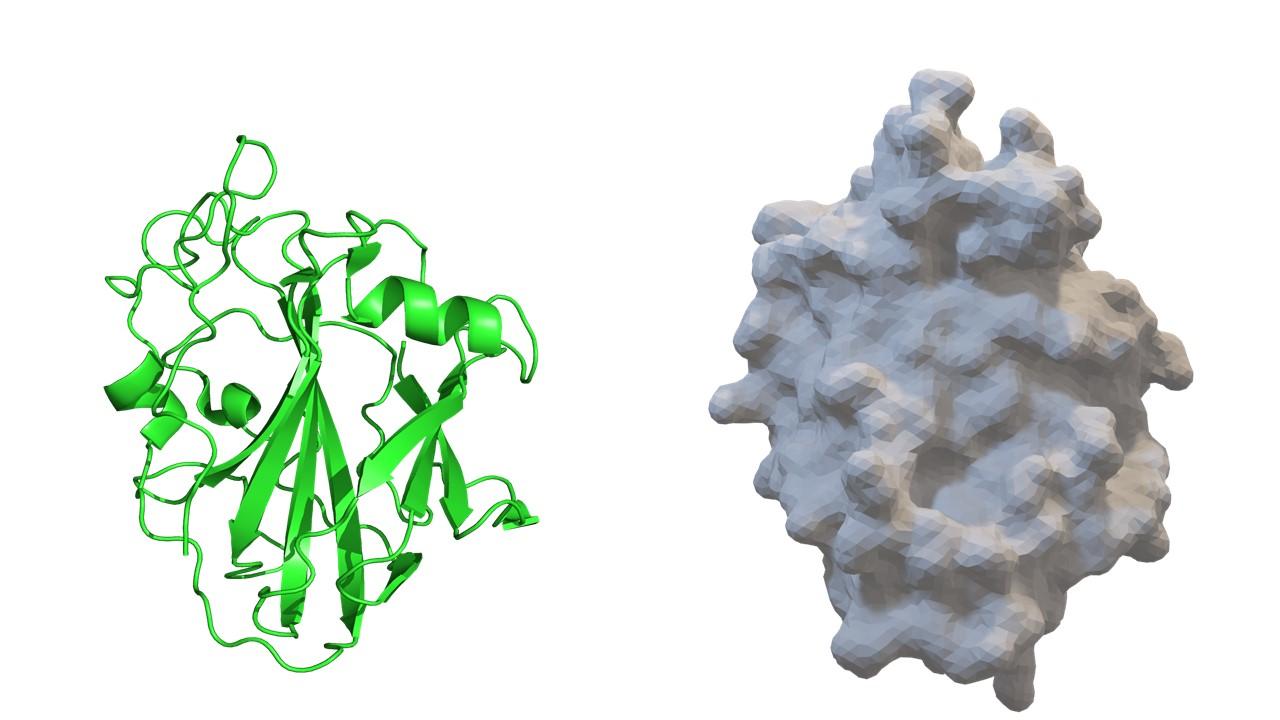

**Supplementary Figure 21: On the left is the molecular representation of chitinolytic LPMO (5NLT) using PyMOL and on the right is the surface representation of a 3D file of the same protein.**
