## Supplementary Tables for "Machine learning-based structural classification of lytic polysaccharide monooxygenases"

**Supplementary Table 1.** Negative and positive data sets were used for the training and validation set considering a ratio of 60:40 split between training and validation sets.

| **Activity of LPMO** | | | | | |
| --- | --- | --- | --- | --- | --- |
| **Positive Set** | | | **Negative Set** | | |
|  | **Number of structures used in Training set** | **Number of structures used in Test set** |  | **Number of structures used in Training set** | **Number of structures used in Test set** |
| **Cellulolytic** | **60** | **40** | **Chitinolytic** | **57** | **38** |

**Supplementary Table 2. Performance metrics of logistic regression using different variants.**

| **Chitinolytic + / Cellulolytic -** | | | | | | | |
| --- | --- | --- | --- | --- | --- | --- | --- |
| **Approach** | | **Logistic Regression** | | | | | |
| **Variant** | | ***lbfgs*** | ***liblinear*** | ***newton-cg*** | ***Newton-Cholesky*** | ***sag*** | ***saga*** |
| **Validation set** | **True Positive** | **32** | **29** | **32** | **32** | **0** | **0** |
|  | **False Positive** | **8** | **11** | **8** | **8** | **40** | **40** |
|  | **False Negative** | **21** | **17** | **21** | **21** | **0** | **0** |
|  | **True Negative** | **17** | **21** | **17** | **17** | **38** | **38** |
|  | **Recall** | **0.63** | **0.64** | **0.63** | **0.63** | **0.49** | **0.49** |
|  | **Precision** | **0.64** | **0.64** | **0.64** | **0.64** | **0.24** | **0.24** |
|  | **F1-score** | **0.62** | **0.64** | **0.62** | **0.62** | **0.32** | **0.32** |
| **Independent Set** | **True Positive** | **1440** | **1443** | **1472** | **1472** | **1448** | **1448** |
|  | **False Positive** | **32** | **29** | **0** | **0** | **24** | **24** |
|  | **False Negative** | **488** | **488** | **517** | **517** | **516** | **516** |
|  | **True Negative** | **29** | **29** | **0** | **0** | **1** | **1** |
|  | **Recall** | **0.74** | **0.74** | **0.74** | **0.74** | **0.73** | **0.73** |
|  | **Precision** | **0.68** | **0.68** | **0.55** | **0.55** | **0.56** | **0.56** |
|  | **F1 score** | **0.65** | **0.65** | **0.63** | **0.63** | **0.62** | **0.62** |

**Supplementary Table 3. Performance metrics obtained for MLP using different solver and activation function.**

| **Chitinolytic + / Cellulolytic -** | | | | | | | |
| --- | --- | --- | --- | --- | --- | --- | --- |
| **Approach** | | **Multi-Layer Perceptron Classifier** | | | | | |
| **Solver** | | ***identity*** | | | ***logistic*** | | |
| **Activation function** | | ***lbfgs*** | ***sgd*** | ***adam*** | ***lbfgs*** | ***sgd*** | ***adam*** |
| **Validation set** | **True Positive** | **32** | **39** | **39** | **40** | **40** | **40** |
|  | **False Positive** | **8** | **1** | **1** | **0** | **0** | **0** |
|  | **False Negative** | **16** | **16** | **16** | **0** | **17** | **1** |
|  | **True Negative** | **22** | **22** | **22** | **38** | **21** | **37** |
|  | **Recall** | **0.69** | **0.78** | **0.78** | **1** | **0.78** | **0.99** |
|  | **Precision** | **0.70** | **0.83** | **0.83** | **1** | **0.85** | **0.99** |
|  | **F1-score** | **0.69** | **0.77** | **0.77** | **1** | **0.71** | **0.99** |
| **Independent set** | **True Positive** | **899** | **1051** | **991** | **954** | **1472** | **983** |
|  | **False Positive** | **573** | **457** | **481** | **518** | **0** | **489** |
|  | **False Negative** | **441** | **453** | **458** | **157** | **517** | **170** |
|  | **True Negative** | **76** | **64** | **67** | **360** | **0** | **347** |
|  | **Recall** | **0.49** | **0.54** | **0.53** | **0.66** | **0.74** | **0.67** |
|  | **Precision** | **0.53** | **0.54** | **0.54** | **0.74** | **0.55** | **0.74** |
|  | **F1 score** | **0.51** | **0.54** | **0.54** | **0.68** | **0.63** | **0.69** |

**Confusion Matrix Components**

**True Positive (TP):** Annotated as cellulolytic LPMO by CAZy and predicted as cellulolytic.

**False Positive (FP):** Annotated as chitinolytic LPMO by CAZy but predicted as cellulolytic.

**True Negative (TN):** Annotated as chitinolytic LPMO by CAZy and predicted as chitinolytic.

**False Negative (FN):** Annotated as cellulolytic LPMO by CAZy but predicted as chitinolytic.
